# Sex change in a protogynous hermaphrodite fish: life-history and social strategies in female cleaner wrasse Labroides dimidiatus

**DOI:** 10.64898/2026.04.06.716686

**Authors:** Letizia Pessina, Redouan Bshary

## Abstract

Protogynous sex change, where individuals first function as females and later as males, is a key life-history strategy among polygynous reef fishes. In haremic systems, sex change is typically socially regulated, with dominants suppressing subordinates’ sex change through aggression. Females within a harem form a size-based hierarchy that can remain stable in most species through the threat of eviction. We studied a different situation in the cleaner wrasse *Labroides dimidiatus*, where larger females have incomplete control, as they spend most of their time alone at their own cleaning territory. We tracked over 400 individuals for 12 months, recording growth, behavior, social organization, and sex change. We confirmed earlier reports that both sexes direct aggression primarily at those ranked immediately below them. However, we observed 30 cases where smaller females outgrew larger ones, revealing hierarchy instability. Of 42 sex change events, 43% occurred in presence of the male, and half of these ‘early’ sex changers were not the largest female, but individuals overlooked by the male. Fast growth relative to harem-mates and harem switching increased the likelihood of sex change. Local population densities also influenced growth and sex change, with individuals in high-density demes growing faster and changing sex at larger sizes. Our findings reveal flexible sex change dynamics in a system with incomplete social dominance. Such incomplete control and observations that becoming male confers both higher reproductive success and survival highlight the need to expand game-theoretical and life-history frameworks to encompass such strategic flexibility.

**Lay summary:** Dominant cleaner wrasse cannot fully control subordinates as individuals occupy distinct core areas. Tracking 400 fish for a year, we found that smaller females could outgrow initially larger ones, and ‘early’ sex change despite a larger male. Fast growth and harem switching increased the chances of becoming male. Population density also shaped these strategies. Our findings reveal flexible sex change dynamics in a system where becoming male confers both higher reproductive success and survival.

## Introduction

Protogyny, a form of sequential hermaphroditism in which individuals first function as females and then as males (Ghiselin 1969; Sadovy and Shapiro 1987; Ross 1990; Munday et al. 2006), is common among fish, particularly wrasses (Kuwamura et al. 2020). Generally, polygynous mating systems, where large males monopolize mating opportunities, favor female-to-male sex change (Munday et al. 2006; Kuwamura et al. 2020). Among protogynous wrasses with known mating system, haremic and lek-like polygyny are the most prevalent (Kuwamura et al. 2020). However, the mating system, the occurrence of sex change, and the proportion of primary (male-born) and secondary (sex changed) males in diandric species can depend on population densities (Warner 1982; Wernerus and Tessari 1991; Kuwamura et al. 2020). These patterns suggest that sex change is not a fixed endpoint, but a flexible strategy through which individuals can maximize their lifetime reproductive success under specific social and ecological circumstances. For instance, males of some fish species adopt alternative reproductive tactics such as early sex change, occurring before the disappearance of the male (Munday et al. 2006), or parasitic tactics, including sneakers and satellites, that allow smaller individuals to exploit the mating opportunities of dominant males (Taborsky 1998; Alonzo et al. 2000; Sato et al. 2004; Oliveira et al. 2008).

Building on this framework, haremic systems offer a particularly interesting context in which to investigate how such flexibility manifests within size-based hierarchies that are presumed to leave little room to alternative strategies. In these systems, a dominant male controls a group of females (Warner and Robertson 1978), and social rank is based on size, with the largest female holding the highest rank below the male (Moyer and Nakazono 1978; Warner 1978; Warner and Robertson 1978; Shapiro 1979; Sadovy and Shapiro 1987; Ross 1990; Devlin and Nagahama 2002; Nelson et al. 2016; Kuwamura et al. 2020). Following the size advantage hypothesis (SAH), which posits that individuals benefit from adopting a sex changing life history when there is a clear difference in the rate at which fitness increases with size for the two sexes (Ghiselin 1969; Warner 1988; Munday et al. 2006), large individuals would optimize their reproductive value by being males that monopolize several females for reproduction (Munday et al. 2006). Importantly, sex change dynamics in a haremic protogynous are primarily governed by males suppressing sex change in their female harem members (Warner 1975; Ross 1990).

The dynamics of a size-based hierarchy and sex change in haremic protogynous hermaphrodites can be understood within the broader context of strategic growth within size-structured groups, where few individuals have privileged access to reproduction. This also includes protandrous sex changers and cooperatively breeding species. Within these systems, conflicts over resources, breeding opportunities, and social rank are most pronounced among individuals of similar size (Enquist et al. 1987; Jennions and Backwell 1996; Cant and Johnstone 2000; Nathan et al. 2001; Bender et al. 2005). This results in selective aggression, where individuals control those directly below them in the hierarchy through aggressive displays in order to maintain the size advantage (Hattori 1991). In return, individuals socially queue and strategically adjust their size and growth rate in relation to those directly above in rank (Hofmann et al. 1999; Kokko and Johnstone 1999; Buston 2003; Heg et al. 2004; Russell et al. 2004; Buston and Cant 2006; Dengler-Crish and Catania 2007; Wong et al. 2007; Young and Bennett 2010; Dubuc and Clutton-Brock 2019), thereby reducing risk of conflict and, in more extreme cases, eviction from the social group (Taborsky 1985; Reeve 1992; Reeve and Nonacs 1997; Reeve et al. 1998; Buston 2003; Buston and Cant 2006; Wong et al. 2007). Similarly, in a protogynous haremic social system, sex change and growth of subordinate individuals are actively suppressed by individuals directly above in rank, with the male primarily focusing on the largest female (Robertson 1972; Moyer and Zaiser 1984; Aldenhoven 1986). This typically results in only the highest-ranking female changing sex at the male’s death, hereafter referred to as standard sex change (Robertson 1972; Robertson 1974a; Robertson 1974b; Warner 1975; Kuwamura 1984; Ross 1990).

Research and conceptual thinking on strategic growth have focused on species that live in cohesive groups. Under these circumstances, subordinates are always in the vicinity of dominants that can hence regularly aggress them, and the threat of eviction is credible. However, in cases of female territoriality, where each female stays in her own preferred core area, while the males try to monopolize the home ranges of several females against other males, harem members are virtually never all together in one cohesive group. This constellation prevents dominants from using eviction as a credible threat. While they may still show aggression toward faster-growing subordinates, they potentially risk losing control, and subordinates might sex change before the disappearance of the male, hereafter referred to as early sex change. The phenomenon of early sex change prompts consideration of potential social dynamics and strategic growth patterns in species with size-based hierarchies if the threat of eviction was absent.

The cleaner wrasse, *Labroides dimidiatus*, provides an ideal model for investigating these dynamics. This species is well known for its complex interspecific social interactions with other reef fishes (Randall 1958; Trivers 1971; Potts 1973; Grutter 1995; Bshary 2001; Grutter 2001; Bshary 2002; Bshary and Grutter 2002a; Bshary and Grutter 2002b; Grutter and Bshary 2003; Bshary and Grutter 2005; Grutter et al. 2005; Bshary and Grutter 2006; Johnstone and Bshary 2007; Pinto et al. 2011), and its social organization has been extensively studied. The cleaner wrasse is a monoandric protogynous hermaphrodite (meaning that all individuals are initially females; there are no initial males) with a size-based hierarchy (Robertson 1972; Robertson 1974a; Robertson 1974b; Kuwamura 1984; Nakashima et al. 2000). Parasitic alternative reproductive tactics are not present as neither males nor females tolerate additional males within their territory (Kuwamura 1984), and spawning occurs exclusively between one male and one female (Robertson and Choat 1974a). Bi-directional sex change has been observed (Kuwamura et al. 2020), but it appears to be entirely governed by social status (Kuwamura et al. 2011). Each female cleaner wrasse maintains her own territory within the range of a dominant male. Female territories can overlap to varying degrees, resulting in either linear or branching social structures, depending on the size differences among females (Robertson 1972; Robertson 1974a; Robertson 1974b; Kuwamura 1984). In branching systems, several groups of females (branches) hold non-overlapping territories, allowing for the coexistence of codominant females, defined as individuals of similar size that avoid interactions with each other (Kuwamura 1984). In contrast, linear systems consist of a single group of females of different sizes with overlapping territories (Kuwamura 1984). Hereafter, we refer to these two configurations as the linear and branching systems. Migration between harems was suggested as a strategy used by lower-ranking females to enhance their opportunities for sex change (Sakai et al. 2001). Classic studies demonstrated sex change suppression and hierarchical control within harems, resulting in standard sex change (Robertson 1972; Robertson 1974a; Kuwamura 1984). Despite these insights, detailed individual-level data on growth trajectories, early sex change, and potential effects of local density are still lacking, leaving the drivers of sex change timing in this species largely unresolved. A focus on individual life histories is of particular interest in light of recent evidence that male cleaner wrasse not only have higher reproductive rates but also higher survival rates compared to females (Pessina et al., in prep.), making sex change a particularly advantageous goal for females.

Here, we conducted a 12-month observational study at Lizard Island, Great Barrier Reef, Australia, to investigate strategic growth and sex change in cleaner wrasse. We marked over 400 cleaner wrasses and collected detailed data on their growth, behavior, and social organization to identify the conditions associated with sex change in individuals.

We use the data to address the following questions. Regarding social control, we ask whether i) we can corroborate earlier results that the hierarchy is largely maintained by each individual keeping the one just below in check; ii) size-based hierarchies are still stable without the threat of eviction, or whether rank reversals occur due to differential growth patterns; and iii) how frequently sex change occurs while the male is still present (compared to after the male disappeared), and whether males fail to exert social control in such cases. Regarding sex change strategies, we ask iv) whether fast growth is a viable strategy to increase chances to change sex and become a male; v) how frequently individuals jump the queue and become a male despite not being the largest female in the harem, and whether this depends on social structure i.e. whether the male has disappeared or is still present, and whether the hierarchy is linear or branching; and vi) whether we can corroborate previous results that switching between harems increases the probability of sex change. Finally, we explicitly compare our study sites and ask vii) whether growth patterns and size at sex change depend on local deme population densities.

## Methods

### Site and Study demes

This study was conducted between July 2022 and September 2023 as part of a long-term observational project at Lizard Island, Great Barrier Reef, Australia. We selected eight reefs (Figure 1) as study demes to capture variation in cleaner and client fish densities.

**Figure 1:**
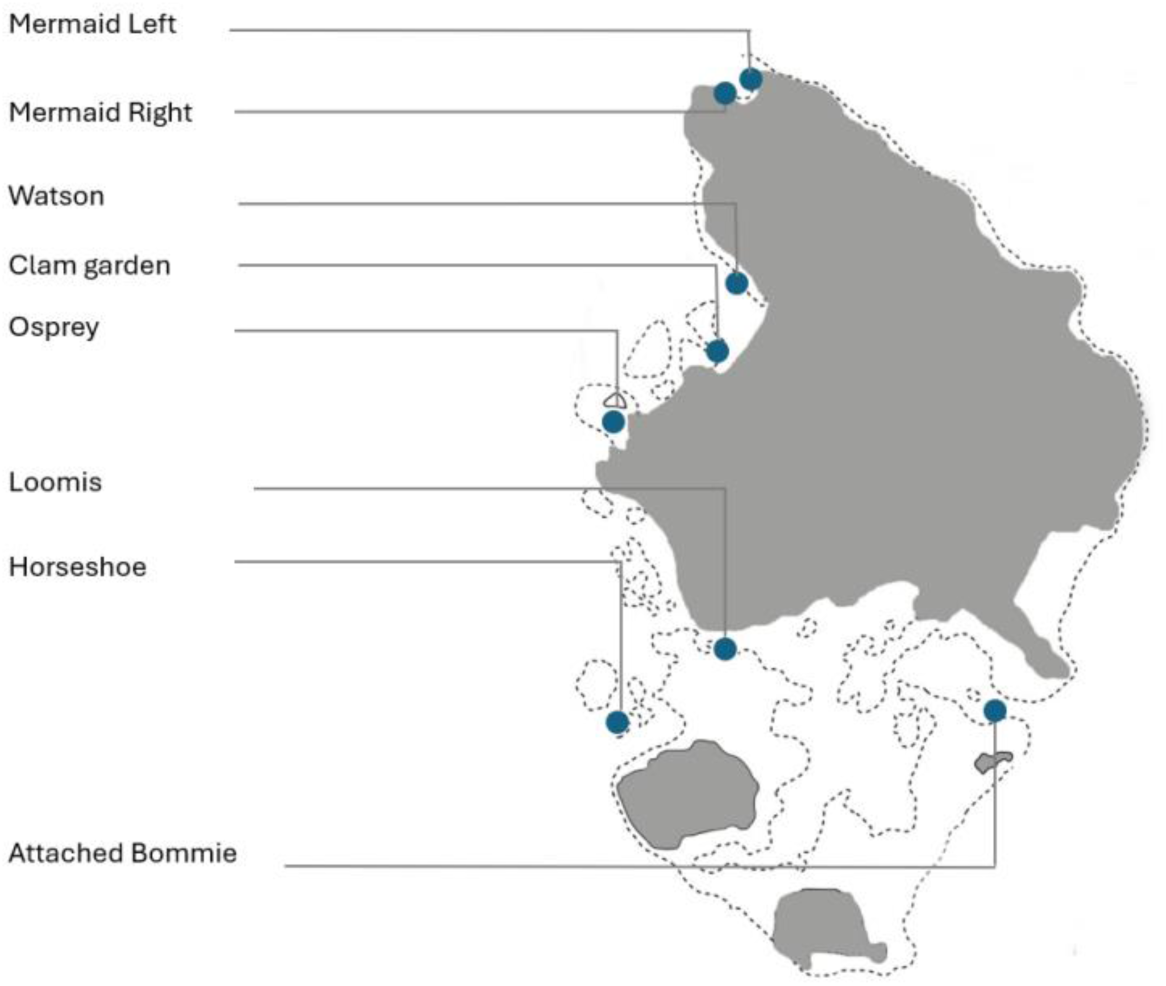
Field observation sites on the coral reefs surrounding Lizard Island, Australia.

Individuals were recognized either by distinctive markings, such as pigmentation patterns or irregularities in the lateral black band (Supplementary Figure S1.2), or by visible implant elastomer (VIE) tags, a common method for small reef fishes (Jungwirth et al. 2019). Tagging was performed underwater after the fish were caught on SCUBA using hand nets (10×15cm) and barrier nets (4.7×1.8m with 2mm mesh for small individuals; 1×1.2m with 5 mm mesh for fish > 6.5 cm), and they were immediately released at the site of capture.

Each adult was marked with two subcutaneous VIE injections placed at distinct locations among four possible body sites (anterior and posterior regions on both sides; Supplementary Figure S1.1). Injections were applied along the pale band above the black stripe using six color options (red, pink, yellow, green, blue, white). Combining two colors across four positions allowed up to 1296 unique identification codes, or more if color order within the same body location was considered (e.g., yellow followed by green or green followed by yellow). Tag combinations could be reused across isolated reefs, as *Labroides dimidiatus* are highly territorial and do not cross open waters. Unique tag sets were maintained only between the two adjacent Mermaid sites.

The VIE pigment-hardener mix (Northwest Marine Technology, Inc. 2017) was applied without hardener to prevent rapid solidification and material loss during large-scale tagging in warm water. Tags remained visible throughout the study despite minor expansion as the fish grew. Recapture of 34 individuals after 12-20 months confirmed the durability of our VIE tags, exceeding previously reported longevity (Jungwirth et al. 2019).

In total, 280 adults were tagged, and 95 were individually recognized by natural features (Supplement Table S1). From November 2022 onwards, approximately 505 juveniles were included in the monitored populations. For the current study, only 165 juveniles that reached a total length of 40 mm are relevant, as they were tagged at that point and subsequently included in the data set. For marking, we used a single contrasting-color injection (red, pink, or yellow), yielding 108 unique tag codes. Overall, 540 individuals were followed, with site-specific sample sizes ranging from 47 (Loomis) to 88 (Mermaid Left).

The primary researcher conducted all fieldwork, totaling ∼1,000 dives and ∼1,750 hours of underwater observation. Continuous monitoring enabled reliable identification of both tagged and untagged individuals through exceptional familiarity with their appearance and behavior.

### Growth

Fish size was measured monthly using a photogrammetric stereo camera system composed of two GoPro Hero 8 cameras (Seager 2006). The system, held by the main diver while swimming through the reef, allowed 3D length measurements from video footage analyzed in EventMeasure (Seager 2006), after calibration in CAL (Seager 2006). Stereo photogrammetry provides higher accuracy than visual estimates (Michael et al. 2011), with reported software errors of 1-2mm (Euan et al. 2010). In our calibration tests, the mean measurement error was ±1.13 mm using a bar of known distances and ±1.81 mm using wild cleaner wrasses, reflecting movement and growth between measurements (See Supplementary Material 2 for full validation details).

When consecutive measurements suggested shrinkage, the previous length was retained as a conservative correction for measurement error. Because individuals were not measured at uniform intervals, we used linear predictions between known size points to estimate the age at which each fish reached successive millimeter increments. This reconstruction produced continuous, evenly spaced size trajectories, ensuring comparability across individuals required to build a size–age growth curve at the demes’ level.

To estimate deme-specific growth rates, we calculated the average time required for fish to grow 1 mm at each site and for each body length. These values provided an expected size-age relationship used to approximate individual ages, as direct age data were not available. Growth curves for each deme were then modeled using the Von Bertalanffy Growth Function (VBGF), fitted via non-linear least squares regression using the nlsLM() function from the *minpack.lm* package (Elzhov et al. 2023).

Distinct deme-level growth trajectories allowed classification of individuals as fast, slow, and average growers at each site. Classification was based on each fish’s final size relative to its deme-specific curve: sizes exceeding half the upper standard deviation limit were labeled fast growers, those below half the lower limit as slow growers, and intermediate values as average growers. The use of half upper and lower limits of standard deviation was chosen to ensure a balanced classification, avoiding an overrepresentation of average growers. Growth-type classification was used to compare survival rates and the occurrence of sex change between growth strategies. To assess the influence of local cleaner densities on the proportions of growth strategies, a secondary dataset classified fast, slow, and average growers relative to a population-level growth curve rather than using deme-specific curves.

### Social system

Sex changing events and key aspects of the cleaner wrasse social structure and dynamics at Lizard Island, including (i) social hierarchies, (ii) formation of linear and branching social systems, (iii) harem fission, and (iv) effects of individual migration, were documented through extensive direct underwater observations. Data were collected bi-weekly via scuba observations and recorded on underwater notebooks.

All individuals within study harems were identifiable through VIE tags or unique morphological features, allowing accurate determination of harem composition, size, and social rank. Changes were tracked by regularly updating individual presence/absence records. An individual was considered absent if no longer seen in the territory, and absence was confirmed after thorough searches in adjacent harems and surrounding reefs. Movements between harems or to peripheral areas were consistently detected and not mistaken for disappearances.

Harem spatial structure was mapped using 2D reef maps generated from DJI Mini 3 Aerial drone footage. Most spatial reconstructions were based on monthly focal videos, with the focal cleaner wrasse’s position plotted every 30 seconds. For some individuals, spatial positions were obtained through direct underwater observations instead. All maps were generated in Adobe Photoshop (Adobe 2019).

Reliable sex identification was essential to describe sex change dynamics. Although cleaner wrasses show limited sexual dimorphism, sex was determined from well-established behavioral cues, which had been validated by Robertson (1974a) through gonadal inspections. This approach allowed long-term, non-invasive monitoring without disrupting social dynamics.

Sex-specific behaviors included: (i) Flutter-run, an exclusively male display involving tail fluttering and fin spreading while swimming past females; (ii) Body-Sigmoid, a female sexual signal where females display an S-shaped posture; (iii) Female-specific sexual coloration during courtship; and (iv) males occupying the superior position during the final phase of the spawning rush. Quantitatively, males showed greater tolerance toward nearby females, higher movement within territories, and more frequent visits to females. Sex classifications were further validated through spawning observations: males spawned only with smaller partners, females only with larger ones.

During the first year, 42 individuals were confirmed to have changed sex, first spawning as females with larger partners and later as males with smaller partners. Consistent with previous findings (Robertson 1974a), following male loss the largest female rapidly displayed male-typical behaviors, signaling an ongoing sex change prior to full physiological transition.

### Intraspecific Behavior observations

Behavioral data were collected for 464 cleaner wrasses across sexes and life stages. Each focal individual was observed at least once per month, yielding 1–13 videos per fish (mean = 4). Among the 42 sex-changing individuals, 0–13 videos were obtained per fish (mean = 7). Only 23 of these were retained for analysis, as others lacked recordings from their female phase.

Each behavioral observation consisted of a 20-minute video filmed with a GoPro Hero 9 by a scuba diver positioned 2-3m from the focal fish. Videos were then analyzed to quantify the number and duration of intraspecific interactions. For each partner, we recorded sex, relative size to the focal individual, and identity (when possible). Absences of interactions were treated as zero values and added during analyses in R Studio (RStudio Team 2020). Social rank for both focal and partner individuals was assigned from the monthly updated social hierarchy dataset.

Behavioral data were used to: i) quantify the time males spent interacting with females before their sex change, and ii) characterize size-based social hierarchies. For the latter, we identified all interactions in which the focal cleaner aggressed their partner through various-intensity chasing behaviors (low: nipping; medium: rushing; high: chase). While preliminary analyses revealed frequent aggression toward small, untagged juveniles without defined rank, only interactions involving individually recognized (tagged or untagged) adults and juveniles with established rank positions were retained. To ensure comparability, we included only focal individuals that could interact with both adjacent and non-adjacent ranks and that had at least 10 video observations. The final dataset included 120 individuals (47 males, 73 females), with behavioral metrics averaged per cleaner.

### Cleaner fish density

We obtained cleaner fish density at our study sites by conducting 10 replicates of 30m transects per site. Counts were stratified into the two habitats of the reef in which our focal cleaner resided: The reef crest and the reef base. The reef crest is defined as the seaward edge of the reef flat, and the reef base as the bottom of the reef slope where it joins the sand flat (Green, 1994). Each fish census involved five 30-meter transects, spaced 2 meters apart, running parallel to the reef edge within each habitat for each site. We conducted counts using a 3m belt, a width previously demonstrated to be effective in counting wrasses (Green 1996). We swam each transect at a constant speed and performed it in approximately 10 minutes. For our analyses, we calculated cleaner densities considering only the counts of adult cleaner wrasse, of which the total length is larger than or equal to 50mm.

### Statistical methods

Data analyses were conducted in R version 4.3.1 (R Core Team 2023) using RStudio (RStudio Team 2020). Depending on the question, we used Generalised Linear Mixed-Effects Models (GLMM), Linear Mixed-Effects Models (LMM), Linear Models (LM), General Linear Models (GLM), and non-parametric tests (e.g., Wilcoxon signed-rank test). Models were fitted using lme4 (Bates et al. 2015), glmmTMB (Mollie et al. 2017), and stats (R Core Team 2023) packages. Model assumptions were verified through residual analyses and visual inspections of model fit. When appropriate, post hoc comparisons were performed with the emmeans package (Lenth 2023). Full model specifications are provided in Supplementary Material 3.

#### 1. Analyses related to Social Control

This section examined whether the cleaner wrasse social system followed a size-based hierarchy and whether dominant individuals exerted effective social control.

To test for hierarchy, we assessed whether aggression was preferentially directed toward the immediately subordinate individual. In each 20-minute focal video, partner ranks were classified as adjacent (directly below the focal individual) or non-adjacent relative to the focal individual. For each video, we calculated the proportion of total aggression directed toward each category and then averaged the values per individual (47 males and 73 females) and across potential numbers of partners. Observed proportions were compared with expected values assuming aggression was evenly distributed across potential partners, using paired Wilcoxon tests on the full dataset and separately for males and females (Models 1-1.3). Videos lacking aggression were excluded from analysis.

After confirming a size-based hierarchy, we assessed the effectiveness of social control by testing whether changes in social rank affected individual growth trajectories. First, we modeled differences in growth rates before and after sex change across different initial sizes using an LMER (Model 2). The log-transformed growth rate was the response variable, and sex, sex change scenario (i.e., standard or early sex change), initial size, and their interactions were included as fixed effects. The female growth rate was calculated using the size measurement 30 days before sex change and at sex change, while the male rates were calculated between the size at sex change and the measurement approximately 30 days after. Accordingly, the initial size refers to the size measured approximately 30 days before sex change for females, and the size at sex change for males. The second LMER model investigates a similar question in the context of females upgrading rank position after the loss of a dominant cleaner (Model 3). Here, Period (Before and After upgrade) was used as a fixed factor, and the log-transformed growth (log(GR 0.3)) was used as the response variable. In both models, Fish ID was included as a random effect to account for repeated measures. Only individuals with both pre- and post-event size data were included (sex change: n = 33; rank upgrade: n = 19). The estimated uncertainty in detecting the exact day of sex change was three to four days, with a maximum of one week in isolated cases.

#### 2. Analyses related to the Social system

We next investigated how features of the social environment influenced the occurrence of early sex change.

Direct field observations determined whether early and standard sex changes occurred more frequently in linear or branching social systems. Two Poisson GLMs were then used to characterize these systems. The first (Model 4) tested whether social structure type, adult cleaner density, and their interaction predicted the number of adult females (TL ≥ 50mm) per harem. The (Model 5) examined whether the likelihood of harem splitting (i.e. the formation of two harems from one) was related to the number of adult females in both linear and branching systems.

A third analysis (Model 6) tested whether reduced control facilitated early sex change. Using an LMER with repeated measures from 23 sex changers recorded within 300 days before sex change, we modeled log-transformed interaction duration (log(duration + 0.3)) as the response variable, with sex change scenario, days to sex change, and their interaction as fixed effects. Fish ID was included as a random factor to account for repeated observations of individuals.

#### 3. Analyses related to Sex change strategies

We then investigated potential strategies that females could use to improve their chances of sex change.

First, we performed two binomial GLM analyses to examine the relation between growth strategy and sex change. The first model (Model 7) tested whether the distribution of fast, average, and slow growers differed between sex-changing (n = 42) and non–sex-changing (n = 334) individuals, including growth strategy, sex change status (sex-changing or not), and their interaction as fixed effects. The second (Model 8) tested whether growth strategy distributions varied among sites differing in adult cleaner reef density, with growth strategy and reef density as fixed factors. Both models were weighted by the number of individuals in each sex change category.

We then explored whether migration influenced size at sex change. The first model (Model 9) tested whether migrants changed sex at a smaller size than residents, using size at sex change as the response variable and migration status (migrant or not), sex change scenario, and their interaction as fixed effects. The second (Model 10) assessed whether size at sex change varied with adult cleaner reef density and the sex change scenario.

## Results

### 1. Social Control

Cleaners directed aggression primarily toward their immediate subordinates rather than lower-ranking individuals (Figure 2, Models 1.1 and 1.2). For both males (Wilcoxon signed rank test with continuity correction: V = 730.5, p-value = 0.038) and females (Wilcoxon signed rank test with continuity correction: V = 2121, p-value < 0.0001), the proportion of agonistic behavior toward adjacent partners was significantly higher than expected under a scenario where aggression was evenly distributed across all lower-ranking individuals.

**Figure 2:**
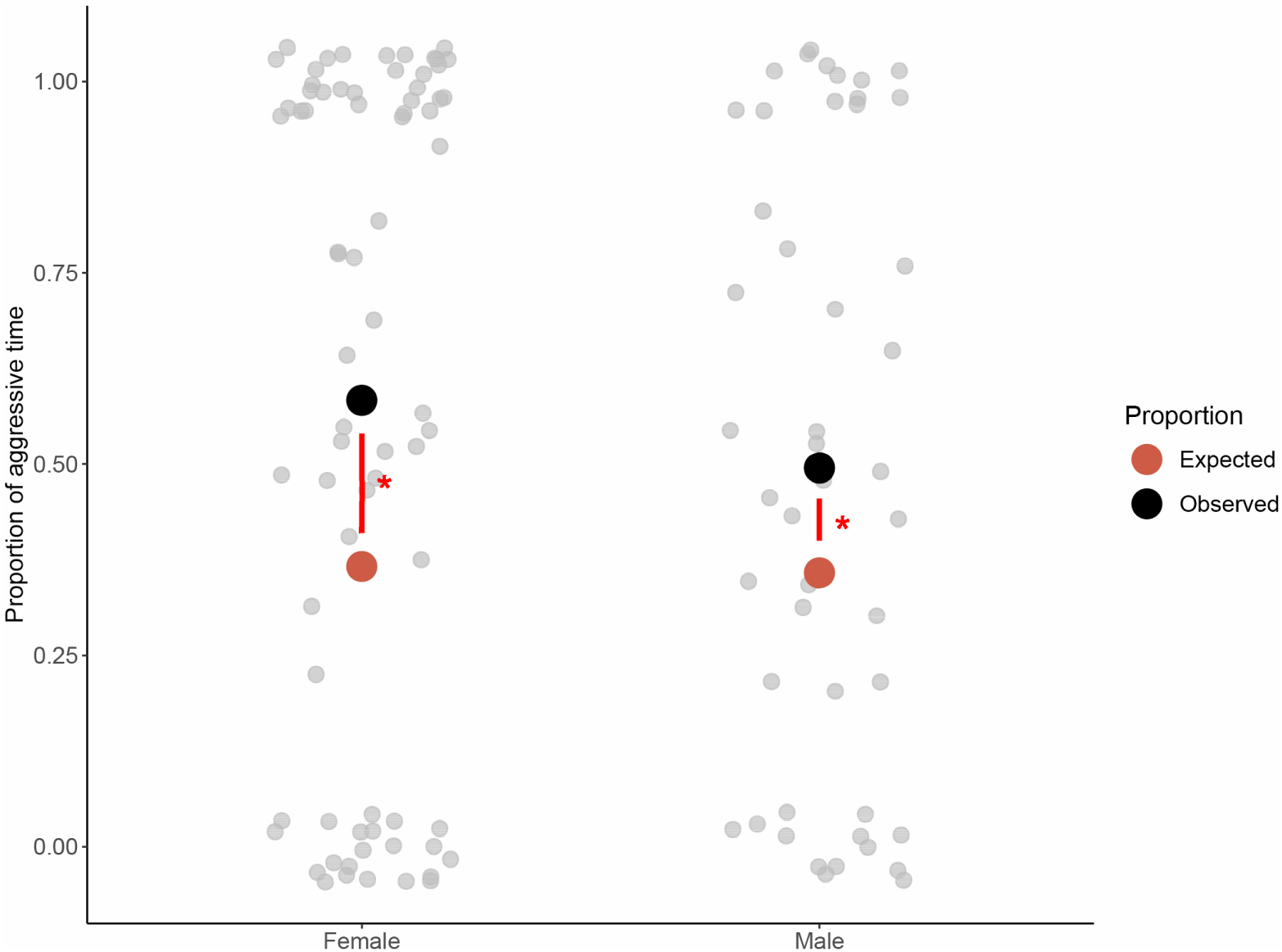
Observed against expected proportion of time spent aggressing adjacent individuals. The distribution of the proportion of time spent aggressing an individual of adjacent rank is shown as a jitter of the raw individual-level observation. Overlayed points show expected (red) and observed (black) mean proportion.

Rank reversals occurred in 20 of 63 harems, with 30 females overtaking the individual directly above them. Growth rates of sex changers significantly decreased after transition (Model 2; Analysis of deviance: Chisq = 4.99, df = 1, p-value = 0.025; Figure 3a), independently of whether it was a standard or early sex change (Analysis of deviance: Chisq = 0.0781, df = 1, p-value = 0.78). Initial size did not significantly interact with the effect of sex change on growth rates (Analysis of deviance: Chisq = 1.25, df = 1, p-value = 0.26), but post hoc tests showed that reduced growth was driven by larger individuals (76–80 mm TL; p < 0.05), whose male-phase growth was 1.5–1.7 times lower than during their female phase (Figure 3a). Conversely, females who moved into higher-quality cleaning stations previously occupied by a higher-ranking female showed a 48% increase in growth following the upgrade (Model 3; Analysis of deviance: Chisq = 6.83, df = 1, p-value = 0.009; Estimate = 0.39, SE = 0.15; Figure 3b).

**Figure 3:**
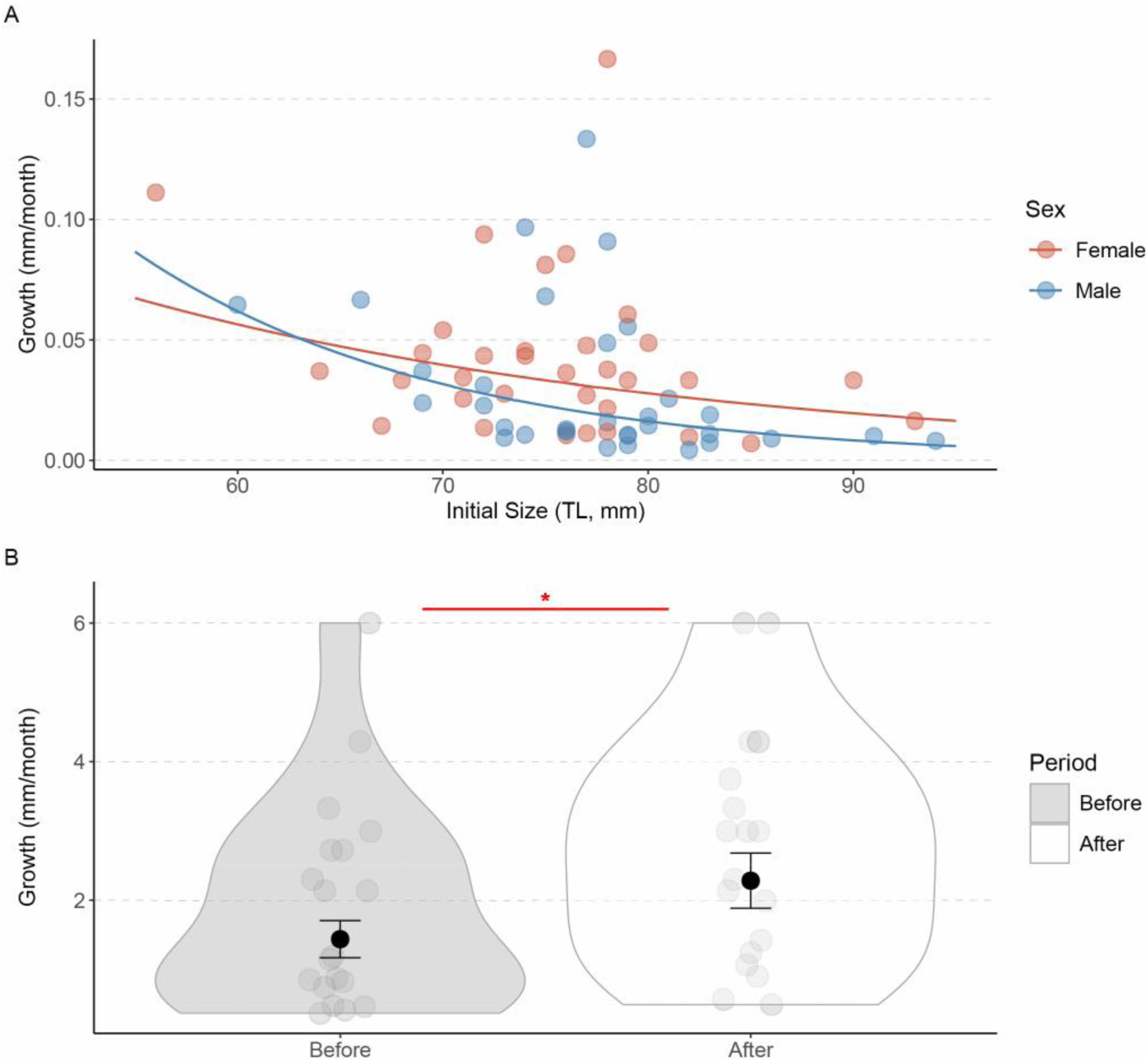
Effects of status upgrade on growth rate. A, Distribution of growth rates (mm/month) across initial body size of the individual before (female) and after sex change (Male), shown as a jitter of the raw observation overlayed by model predictions. B. Distribution of growth rates (mm/month) before and after upgrading rank position as a female, shown as violins and jitter of the raw individual level observations. The overlaid points and error bar represent model-estimated means ± standard errors (SE).

### 2. Social System

At the beginning of the study, the adult sex ratio was 4.2 females per male; by the end, it was 4.4. Among the 196 initial females, 42 underwent sex change between July 2022 and September 2023. Of these, 57.1% (n = 24) and 42.9% (n = 18) were standard and early sex changes, respectively. 88.5% of the standard sex changers were the largest and most dominant females in their harems. In contrast, only half of the early sex changers were dominant, with the remainder being smaller, lower-ranking females.

Early sex change was closely linked to the social structure. In particular, while the majority of standard sex changes occurred in linear social systems (88.3%), most early sex changes occurred in branching social systems (77.8%). Branching systems were always associated with the presence of codominant females, defined as large individuals of similar size in the male’s territory. Codominant females were not present in any of the linear systems (see Supplementary tables S14.1.1 and S14.2.1).

In the few cases of standard sex change that occurred in a branching system (16.7%, nb = 4), pairs of codominant females changed sex at the same time and split the original harem (see S14.2l, S14.2o). Three of the early sex changers that belonged to a linear system were individuals that migrated at the periphery of the male’s territory and created their own harem instead (see S14.1h, S14.1f). The remaining one was abandoned by its male (see S14.1l).

Harems of more adult females (≥ 50mm) were significantly more likely to form branching systems than linear systems (Model 4, Analysis of Deviance: LR Chisq = 25.35, df =1, p-value <0.0001, Figure 4a) and were also more likely to split and create two or more new harems (Model 5, Analysis of deviance: Chisq = 24.00, df = 1, P-value < 0.0001 ; Figure 4b). Adult cleaner reef density did not significantly affect adult female harem size (Analysis of deviance Type II: Chisq = 1.13, df = 1, P-value = 0.29).

**Figure 4:**
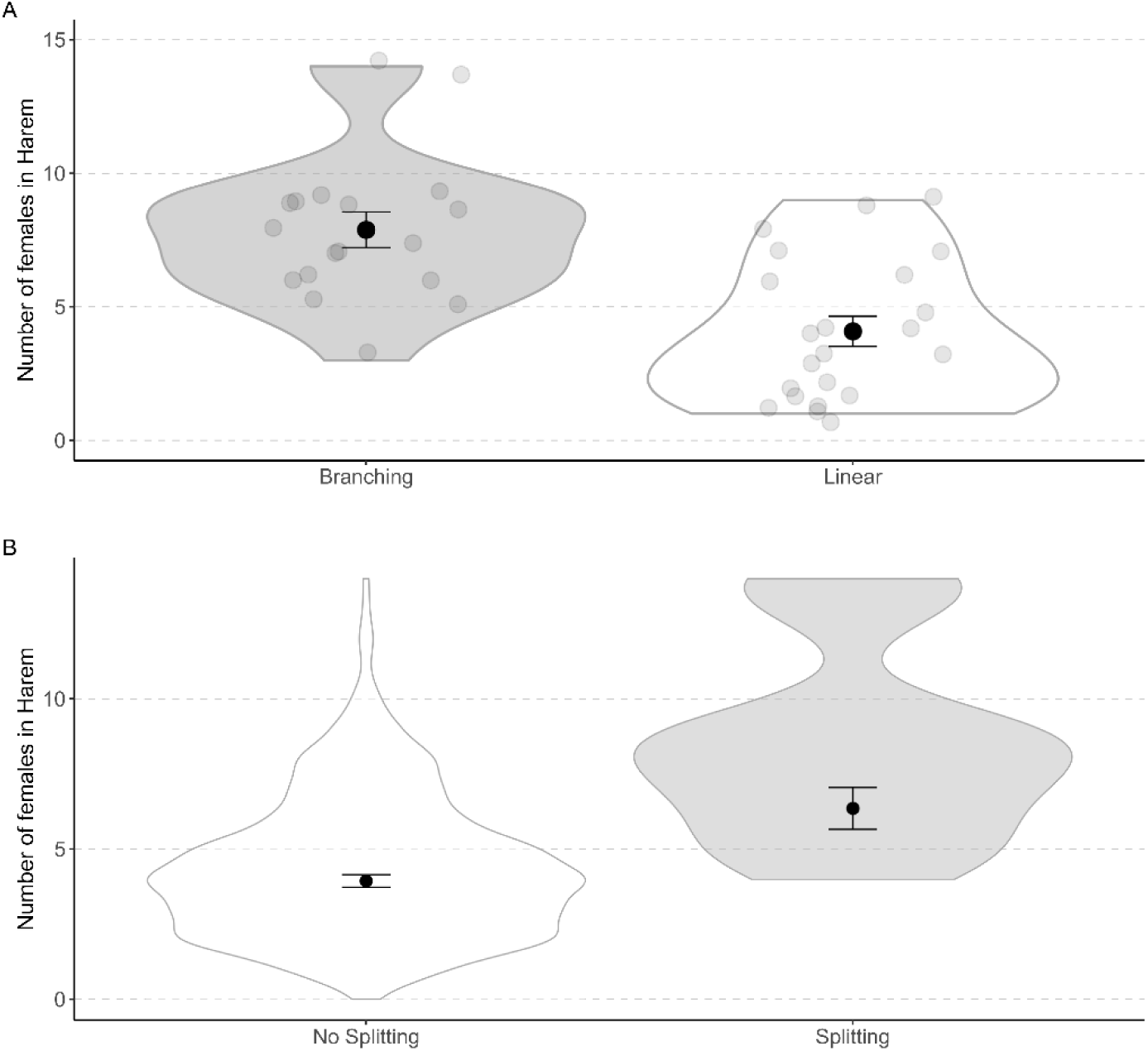
Number of adult female cleaner wrasse in various scenarios. Distribution of the number of adult females (TLTL ≥ 50mm) within branching and linear social systems (A), and within harems that split into two or that did not split (B), shown as violins and jitter of the raw transect-level observations at each study site. The overlaid points and error bars represent model-estimated means ± standard errors (SE).

Analyses of male–female interactions before sex change, revealed that males spent significantly more time with standard sex changers than with early sex changers (Model 6; Analysis of deviance: Chisq = 7.92, df = 1, p-value = 0.005). Interaction time increased significantly as females approached sex change (Analysis of deviance: Chisq = 15.24, df = 1, p-value < 0.001). This was true for both standard and early sex changers (Analysis of deviance: Chisq = 0.014, df = 1, p-value = 0.90; Figure 5). Post hoc comparisons indicated that standard sex changers (emmeans = 1.088, SE = 0.71, on the log-scale) spent almost nine times longer interacting with males than early sex changers (emmeans = 3.16, SE = 0.69, on the log-scale).

**Figure 5:**
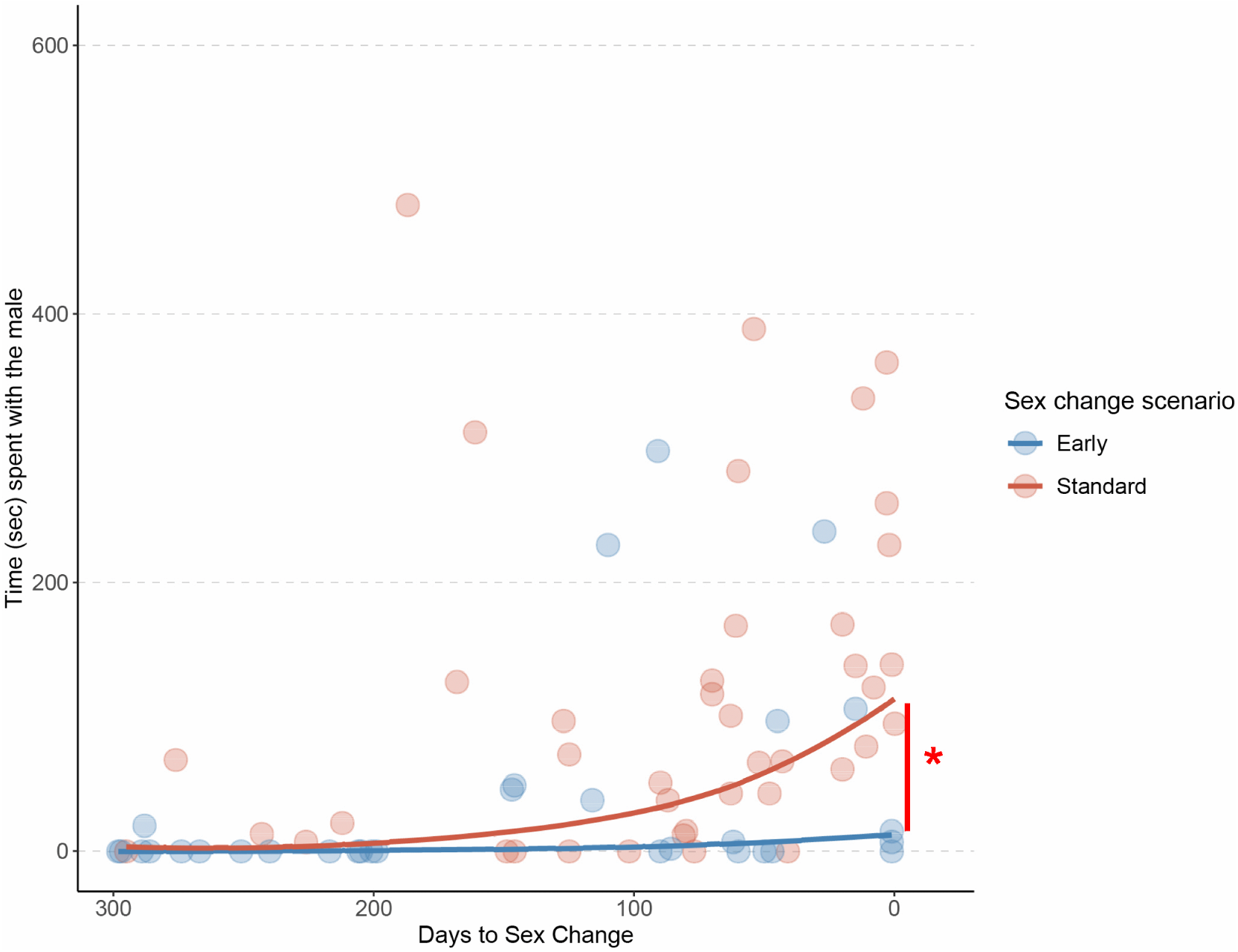
Time males spent with sex-changing females prior to sex change for the two sex change scenarios. Distribution of the time males spent with the sex-changing female during 20-minute video observations taken at different moments prior to their sex change for each sex change scenario. Raw individual observations are shown as jitter, and the overlaid lines represent model predictions.

### 3. Sex Change Strategies

Fast growth emerged as a key intrinsic factor facilitating sex change. Although the average growth strategy was most common across the population (Figure 6a), growth strategy distribution differed significantly between sex-changing and non-sex-changing individuals (Model 7; Analysis of deviance: LR Chisq = 18.17, df = 2, p-value < 0.001). Post hoc comparisons showed that fast growers were 87.5% more frequent among sex changers (45%) than among non-sex changers (24%; Estimate = -0.972, SE = 0.335, p-value = 0.0038), whereas slow growers were significantly underrepresented (14% vs. 38%; Estimate = 1.281, SE = 0.455, p-value = 0.0049) Average growers showed no difference between the two groups (Estimate = -0.075, SE = 0.334, p-value = 0.8229).

**Figure 6:**
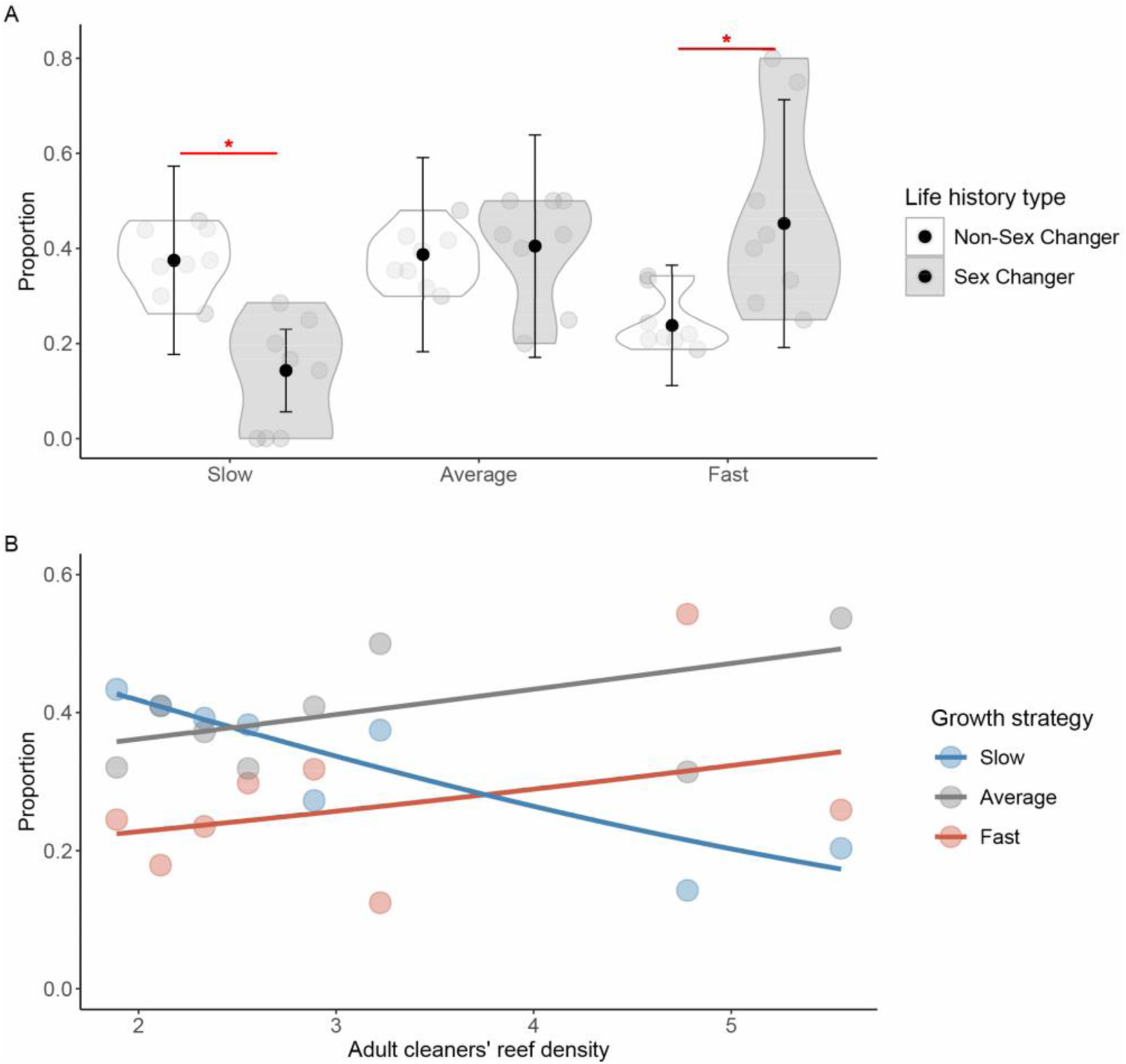
Proportions of fast, slow, and average growers. A. Distribution of the proportions of fast, slow, and average growers within sex and non-sex changers at each site, shown as violins and jitter of the raw site-level observations. The overlaid points and error bar represent model-estimated means ± standard errors (SE). B. Distribution of the proportion of fast, slow, and average growers across different adult cleaner wrasse reef densities (TL) shown as a jitter of the site-level raw observation overlaid by model predictions.

Pooling all data to calculate an overall population growth curve revealed that the distribution of growth strategies varied with adult cleaner density (Model 8, Analysis of deviance: LR Chisq = 19.79, df = 2, p = 0.0001; Figure 6b). Both fast and average growers increased in frequency with density (Average trend: estimate = 0.151, SE = 0.123; Fast trend: estimate = 0.161, SE = 0.09), while slow growers declined (trend estimate = -0.346, SE = 0.996).

Across sites, the mean size at sex change was 78.35mm (TL). Size at sex change increased significantly with adult cleaner reef density (Model 10, Analysis of deviance: F = 9.35, df = 1,p-value = 0.004; Figure 7), while sex change scenario (standard vs. early) had no effect on size at sex change (Analysis of deviance: F = 6.71,df = 1, p-value = 0.66; Figure 7).

**Figure 7.**
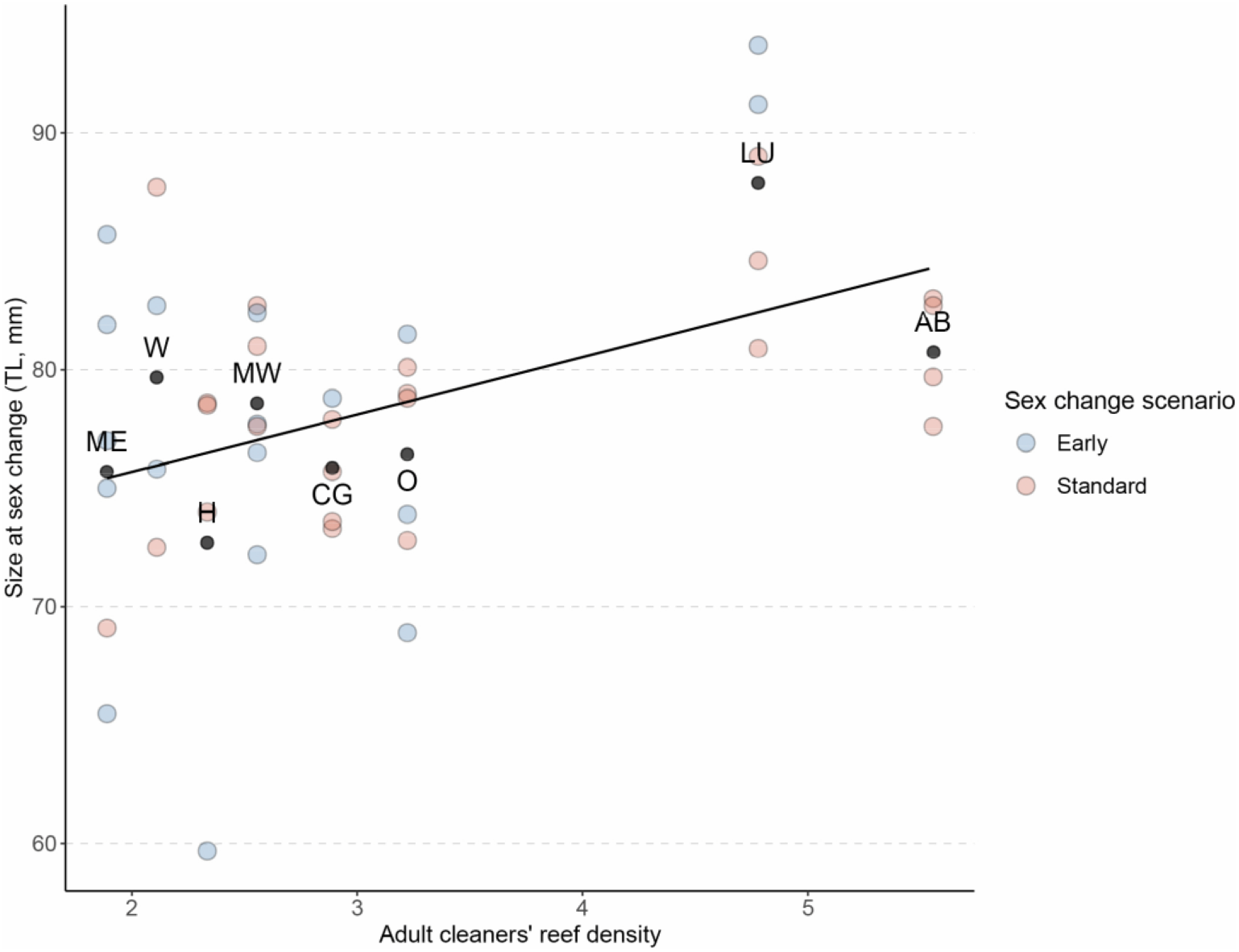
Size at sex change across adult reef cleaner densities. Distribution of size at sex change (TL, mm) plotted against adult cleaner fish reef density, shown as a jitter of the raw individual-level observations at each site overlaid by the predictions of the model.

Migration to neighboring harems also influenced sex change. Individuals that migrated before transitioning changed sex at significantly smaller sizes than non-migrants (Model 9; Analysis of deviance: F = 5.33, df = 1, p-value = 0.027; Figure 8a), with no interaction between migration and sex change scenario (Analysis of deviance: F = 0.95, df = 1, p-value = 0.33; Figure 8a). On average, migrants changed sex at 6% smaller body size than non-migrants (early sex change: 73.9 ± 2.23 mm vs. 80.7 ± 1.99 mm; Standard sex change: 76.6 ± 2.23 mm vs. 79.5 ± 1.58 mm).

**Figure 8:**
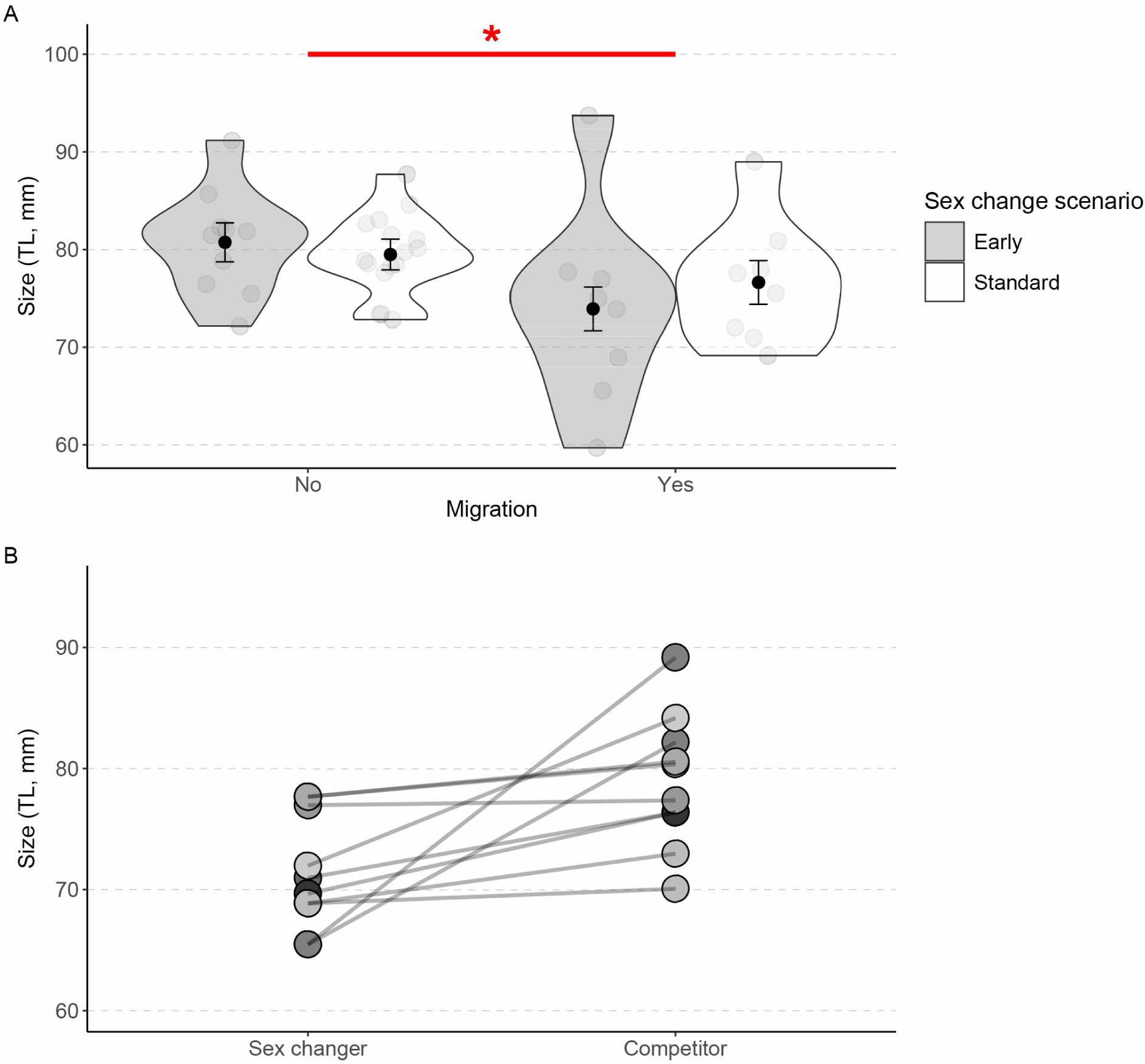
Effects of migrating. A. Distribution of size at sex change (TL, mm) for migrant and non-migrant females in the two sex change scenarios shown as a jitter and violins of the raw individual-level observations. The overlaid points and error bar represent model-estimated means ± standard errors (SE). B. Size at sex change (TL, mm) of seven migrant females compared to the size of their competitors in the original harem at the time of sex change.

Overall, 38,1% (16 of 42) of the sex changers were migrants at a certain point in their life, compared with a 12.7% migration rate in the general population (37 of 291), corresponding to a 43.2% probability of sex change in migrant versus 10.2% in non-migrant female population.

Excluding the four individuals for whom we have no information before their migration, 91.7% (11 of 12) migrant sex changers were low-ranking females in their previous harem. For seven migrant sex changers, we have additional information about their original harem (Figure 8b). All of them would not have had the opportunity of changing sex in their original harem at the time of their sex change, as they would still be occupying a low rank position (either Beta or lower, Figure 8b).

Most migrants transitioned within linear systems (68.75%). The number of migrant standard sex changers (n = 8) was similar to the number of migrant early sex changers (n = 7). Of the 8 migrant early sex changers, four transitioned after the male abandoned a branch within a branching system, one after the male left his linear harem, and three after moving to peripheral territories to establish new branches.

Six individuals changed sex at sizes smaller than 75mm, raising the question whether sex change was beneficial or rather the product of a mechanism (‘change sex if not regularly visited by a male’) wrongly triggered. Three of these small individuals migrated to the periphery of their social group, and formed new harems with only one to three juveniles, whereas the remaining three gained adult females as subordinates. The 10 larger sex changing females (≥ 75 mm) typically obtained one to five adult females and up to four juveniles after transitioning (see Supplementary Material 15).

## Discussion

Our study builds on previous research on the cleaner wrasse social system to understand strategic growth decisions in a protogynous fish species where dominants lack the threat of eviction as a control mechanism to suppress fast growth or sex change. We confirmed various previous results, like i) the coexistence of linear and branching hierarchies within a harem; ii) that individuals of both sexes mainly aggress the harem member that is below them in the hierarchy as a means to maintain dominance; and iii) that switching between harems may be a viable strategy to improve one’s own rank. Our additional data yield new insights into strategic growth as a function of a rise in hierarchy, social structure, and population density, which in turn affects the probability that an individual changes sex and becomes male. We discuss these results in detail below.

### 1. A Size-based hierarchy with incomplete control

The cleaner wrasse at Lizard Island displayed higher levels of aggression towards adjacent individuals compared to what would be expected if they were to aggress at each rank below in the same manner. This finding, which agrees with previous results (Kuwamura 1984), supports a key characteristic of a size-based hierarchy, as in such systems, aggression is typically highest between individuals of neighboring ranks (Robertson 1972; Moyer and Zaiser 1984; Aldenhoven 1986). Nevertheless, the size-based hierarchy does not translate into the strategic growth patterns that have been described for species in which the dominant may evict subordinates that grow too fast. According to existing strategic growth theory and data, individuals in a size-based hierarchy are expected to accelerate growth when individuals above them in rank disappear (Hofmann et al. 1999; Kokko and Johnstone 1999; Buston 2003; Heg et al. 2004; Russell et al. 2004; Dengler-Crish and Catania 2007; Wong et al. 2007; Young and Bennett 2010). While females showed increased growth when a higher-ranking individual disappeared, this seems to be linked to them occupying the cleaning station of the disappeared individual. This switching was a general feature, suggesting that one benefit of climbing in the hierarchy is to be able to occupy a station that offers a better client composition and hence more food. This aspect may ‘naturally’ stabilize the social hierarchy. However, we documented various rank reversals due to initially smaller individuals outgrowing initially larger ones. As far as we are aware, such cases have not been observed in species with the threat of eviction. Finally, sex change and resulting top dominance position did not lead to faster growth but instead to reduced growth.

Another line of evidence for the incomplete control in the size-based hierarchy of the cleaner wrasse at Lizard Island is that early sex change occurs regularly. 42.9% of the sex-changing events happened in the presence of the male. Previous research has already documented the occurrence of early sex change in this species (Robertson 1972; Robertson 1974a; Kuwamura 1984). However, since most of the research was based on experimental removals of males, the magnitude of its occurrence was masked by the manipulations. This study, therefore, brings novel evidence of the limited control that a male cleaner wrasse has over the sex changing tendencies of some of his females at Lizard Island. In addition, dominant females also have incomplete control, as we documented sex change by individuals that were not the largest female in the harem. Such early sex changes typically occurred in branching systems, in which the largest female would still be aggressed by the male but would not interact with the next largest. Here, referring to the game theory model of the three-person duel, the “weakest” female prevails over the “strongest” (Archetti 2012; Dorraki et al. 2019). The results challenge current theories for which in a protogynous hermaphrodite with sex change suppression, early sex change should not be as common (Ross, 1990; Warner, 1975). Our new results are particularly relevant as *Labroides dimidiatus* is the classic example of sex change suppression (Warner 1975; Ross 1990).

### 2. Individual sex change strategies

Given the apparent instability of the size-based hierarchy, one could expect that some features of individuals and opportunities increase or decrease the likelihood of sex change. One such feature we identified was speed of growth; fast growers were most likely to change sex. In principle, fast growers could be individuals of high quality (either as successful cleaners or because of efficient immune function) or they might trade off current reproductive success (egg production) against potential future reproductive success. However, recent evidence that male cleaner wrasse have higher survival rates than females do (Pessina et al., in prep.), on top of the higher reproductive rate, suggests that any decisions that increase the likelihood of eventually achieving sex change are the best possible options, and that only low-quality individuals would opt out of the general competition and focus on current reproductive success.

Migration to a more suitable harem also emerged as a strategy used by females to improve their chances of sex change. This result aligns with previous observations (Sakai et al., 2001). As harem compositions are outside the control of individuals, switching will always be an opportunistic decision. At this stage, we do not know whether there is inter-individual variation in prospecting neighboring harems or whether all individuals have knowledge about the composition of neighboring harems.

As an important addition to the existing literature on cleaner wrasse, our results suggest that sex changing at a smaller size may not always be advantageous. Individuals that started their harem by having only juvenile harem members reduced their reproductive rate compared to the alternative of having remained a female and have own eggs fertilized by the male harem owner. Early laboratory experiments on cleaner wrasse suggest that the social control mechanism responsible for sex change suppression may sometimes cause wrong decisions: medium-sized females can be induced to change sex when taken away from the male and paired with a smaller female, but then may reverse back if paired again with a larger male (Kuwamura et al. 2002; Kuwamura et al. 2011).

### 3. Individual strategies as a function of local densities

Somewhat surprisingly to us, the number of females in a harem and the social dominance structure (linear versus branching) were not affected by cleaner density. Thus, the documented positive correlation between population density and size at the moment of sex change cannot be explained by increased competition within a harem. Instead, the alpha female’s potential to sex change may be threatened by neighboring individuals who could take over the harem if she is not sufficiently large to compete with them following the disappearance of the male (Robertson 1972). Alternatively, as cleaner density is known to correlate with the density of large client fish (Triki et al. 2019), individuals growing up in high-density sites might simply grow faster because of access to more food resources and shift partly the tradeoff between current egg production and investment in becoming a male. Both potential explanations fit the observed positive correlation between local population density and the proportion of fast-growing individuals. Local population density is hence an important factor affecting growth and sex change decisions in cleaner wrasse. Its importance for strategic growth in species living in cohesive groups with subordinates facing the threat of being evicted remains to be investigated.

## Conclusion

In conclusion, our study on cleaner wrasse yields important new insights into social hierarchies, strategic growth, and sex change decisions in a species in which dominants have incomplete control due to the lack of group cohesiveness and in which becoming a male confers both higher reproductive success and survival. The new phenomena that we describe, relative size reversal, early sex change not being restricted to the largest harem female, effects of population density on growth and sex change patterns, reveal the need for a new generation of life history models that vary the degree of dominant control in such social systems.

## Acknowledgements

We thank the staff of the Lizard Island Research Station in Australia, L. Vail, A. Hoggett, R. Carr, and A. Davie, for their exceptional support and for fostering an outstanding research environment. We are grateful to Dr. F. Cortesi and the University of Queensland for hosting, and to Dr. A. Green for training in transect data collection. We also thank A. Viglino, M. Amann, S. Deventer, E. Pattenden, S. Lévy, C. Lebet, and D. Berger for their invaluable assistance in the field, and Dr. Radu Slobodeanu for statistical guidance. A special thanks to the inspiring young researchers we met on the island, whose enthusiasm and thoughtful conversations made the work environment both stimulating and welcoming. This work was supported by the Swiss National Science Foundation (grant 310030_192673/1 to RB).

