## Supplementary Materials for "Sex change in a protogynous hermaphrodite fish: life-history and social strategies in female cleaner wrasse Labroides dimidiatus"

### Supplement 1: Specific numbers of cleaner wrasse per site

#### Table S1: Total number of focal cleaner wrasses at the 8 sites.

This table summarizes he total number of tagged (VIE) and non-tagged (No VIE) for the 8 sites: Mermaid Left (ML), Mermaid Right (MR), Watson (W), Clam Garden (CG), Osprey (O), Horseshoe (H), Loomis (LU), and Attached Bommie (MB).

| Site | Adult Males | | Adult Females | | Juveniles | Total | Sex changers |
|---|---|---|---|---|---|---|---|
| | VIE | No VIE | VIE | No VIE | VIE | | |
| **ML** | 3 | 1 | 45 | 5 | 34 | 88 | 6 |
| **MR** | 6 | 0 | 29 | 10 | 28 | 73 | 7 |
| **W** | 3 | 3 | 31 | 8 | 19 | 64 | 4 |
| **CG** | 5 | 1 | 31 | 9 | 12 | 58 | 5 |
| **O** | 2 | 3 | 33 | 10 | 19 | 67 | 7 |
| **H** | 6 | 0 | 38 | 17 | 18 | 79 | 4 |
| **LU** | 3 | 1 | 18 | 9 | 16 | 47 | 5 |
| **MB** | 5 | 5 | 22 | 13 | 19 | 64 | 4 |
| | 33 | 14 | 247 | 81 | 165 | 540 | 42 |
| **Total** | 47 | | 328 | | 165 | 540 | 42 |

#### Figure S1.1: Illustration of the four locations for VIE

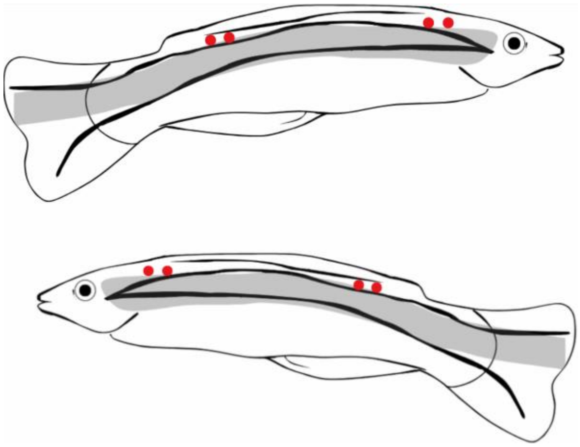

**Figure S1.2: Examples of two individuals that can be recognized without VIE**

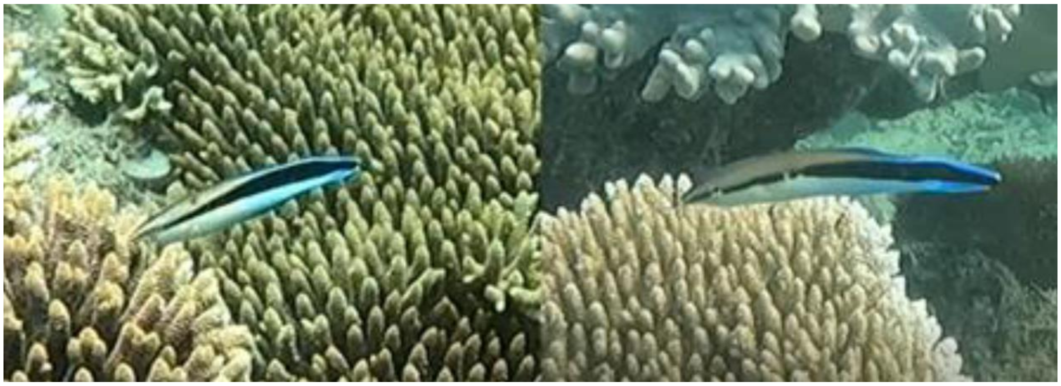

### Supplement 2: Stereo camera sizing error

We measured the size of each focal fish every month, using two GoPro Hero 8 cameras mounted in a photogrammetric underwater stereo camera system (Seager, 2006). Sizing footage was analyzed using Event Measure (Seager, 2006), a software enabling 3d length measurements. This method relies on prior calibration of the stereo camera system performed by the CAL software (Seager, 2006). Utilizing a stereo camera system enhances accuracy and precision compared to visually estimating fish size (Michael et al., 2011). The error of this software is known to be around 1-2mm (Euan et al., 2010). To further assess the accuracy of the stereo camera system for our study, a calibration bar with three known distances was used to measure the system's error. Measurement error was ± 1.13mm using the calibration bar and ± 1.81mm of wild cleaner fish due to movement and growth between measurements (Figure S2, Table S2).

The overall mean error obtained with the toolbar was ± 1.13mm. Because measuring a moving object is more challenging, the software's accuracy was then investigated by comparing the stereo camera measurements of fish longer than 70cm with manual size measurements taken less than 30 days prior. This method provided an average error of ± 1.81mm. Attempts were made to obtain size measurements using the stereo system within 1 week of the fish being manually measured post-capture. However, this proved challenging, as the fish required more time to re-acclimate to human presence and often swam too quickly or attempted to escape, making accurate measurements difficult within this short time frame. Nevertheless, the errors associated with our fish measurements did not show a significant increase in variance (variance = 2.324) compared to those observed with the calibration toolbar (variance = 2.264). This suggests that the software's measurement error remains consistent when applied to real fish. The positive shift in the median error for the fish measurements (median = 1.48 mm) reflects the fish's natural growth over the 30 days between the two measurements. This consistency in error variance across both methods indicates that the software performs reliably for measuring wild fish.

### Figure S2: Sizing error using the calibration tool and the focal cleaners

The following figure shows the size error of the software obtained using sizing focal cleaners (“Cleanerfish”) measured 30 days from their manual sizing, and using the objects of known size (“Calibration”).

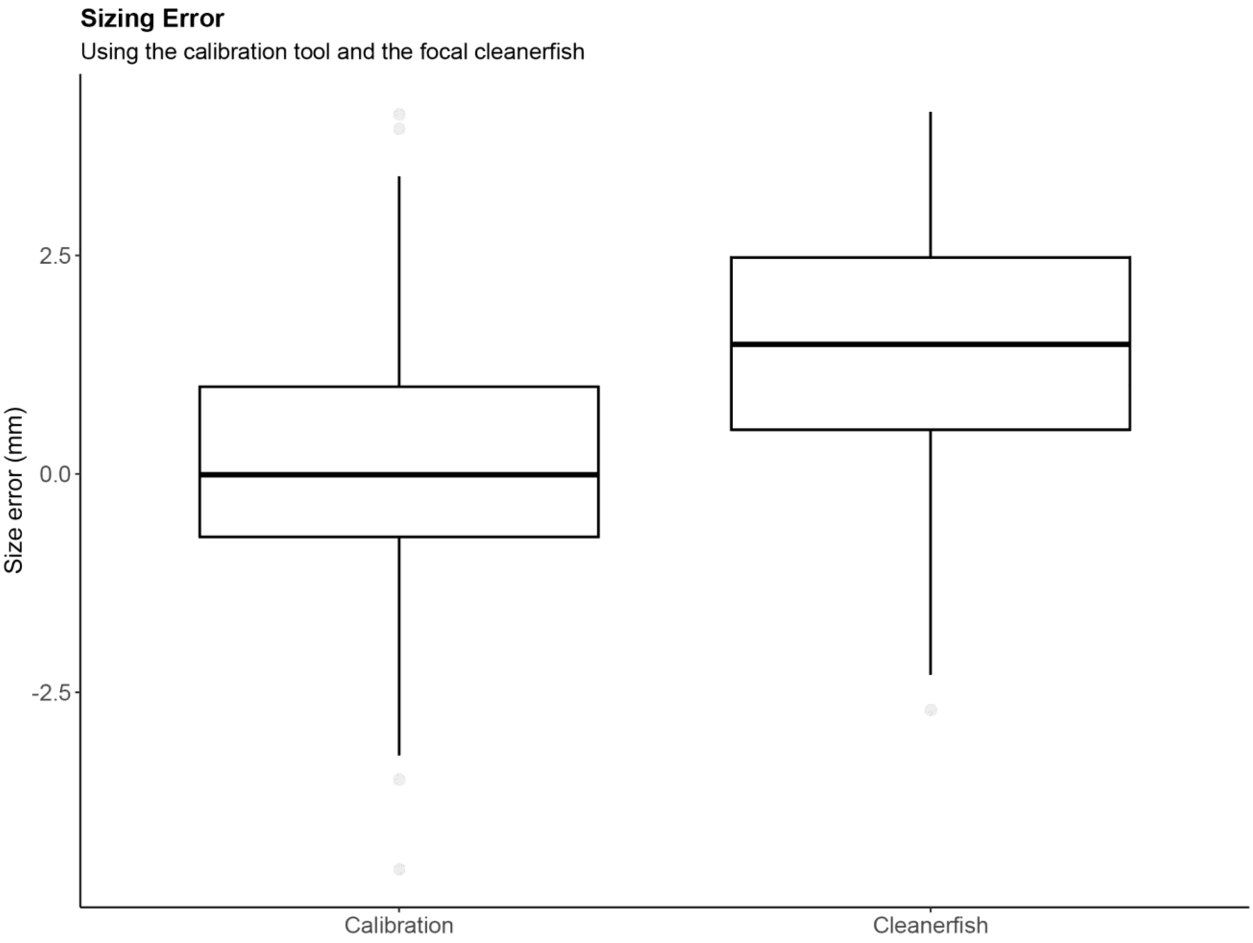

**Table S2: System’s errors obtained with the different methods:**

| Method | Mean Error (mm) |
|---|---|
| Small | ± 0.981 |
| Medium | ± 1.08 |
| Large | ± 1.37 |
| General Tool | ± 1.13 |
| Fish | ± 1.81 |

### Supplement 3: Model Details and Assumption Checks

**Table S3.1: Glossary for tables 4.2 and 4.3**

| Term | Definition |
|---|---|
| Days | Days to sex change |
| Density | Adult cleaner fish reef density within a deme (per $150m^2$) |
| Females | Number of adult females in a Harem |
| GR | Growth rate (mm/month) |
| ID | Cleaner ID used as a random factor to control for repeated measures |
| Interaction | Total time spent with the male (in seconds over a 20-minute video) |
| Initial Size | The size measured approximately 30 days before sex change for the female phase, and the size at sex change for the male phase |
| LifeHistory | Sex changer or non-sex changer |
| Migration | If the fish migrated to another harem (yes or no) before their sex change |
| Period | Before and after upgrading the cleaning station |
| Prop | Proportion |
| Scenario | Early or standard sex change |
| Size | Size of the cleaner fish (TL in mm) |

### Table S3.2: Model Description

| Model | Description |
|---|---|
| 1 | Tests the differences between observed and expected proportions of aggression directed toward adjacent individuals. |
| 1.1 | Tests differences between observed and expected proportions of aggression directed toward adjacent individuals, taking only males into account. |
| 1.2 | Tests differences between observed and expected proportions of aggression directed toward adjacent individuals, accounting only for females (Alpha, Beta, and Gamma). |
| 2 | Examines the effect of sex change on growth rate. |
| 3 | Examines the effect of rank and cleaning station upgrade on growth rate. |
| 4 | Investigates predictors of branching versus linear social systems. |
| 5 | Investigates the relationship between the number of adult females in a harem and the likelihood of a splitting event (i.e., an harem splits into two) in both Systems. |
| 6 | Investigate whether sex change in the presence of the male is associated with a reduction of social interactions (duration in seconds) with the latter. |
| 7 | Tests whether a specific growth strategy increases the likelihood of sex change. |
| 8 | Examines whether reef density influences the proportions of different growth strategies. |
| 9 | Investigate the size at sex change in migrants versus non-migrants across sex change scenarios. |
| 10 | Assesses how reef cleaner density and sex change scenario affect size at sex change. |

### Table S3.3: Model Summary

| Model | Type | Family(link) | Challenge | Formula | Sample Size |
|---|---|---|---|---|---|
| **1** | Wilcoxon test | - | Non-normality | Prop, expected, paired = T | N = 120; Cleaners = 111 |
| **1.1** | Wilcoxon test | - | Non-normality | Prop, expected, paired = T | N = 47, Cleaners= 47 |
| **1.2** | Wilcoxon test | - | Non-normality | Prop, expected, paired = T | N = 73, Cleaners = 69 |
| **2** | LMER | Gaussian | Right Skew | log(GR)~Sex + Scenario + initial-Size+ Sex:Secnario + Sex:Initial_Size + (1\|ID) | N = 66; Cleaners = 33 |
| **3** | LMER | Gaussian | Right skew | Log(GR+0.3)~Period+(1\|ID) | N = 38; Cleaners = 19 |
| **4** | GLM | Poisson (log) | - | Females~System*Density | N=40 |
| **5** | GLMM | Poisson (log) | - | Females ~Splitting + (1\|Site/Harem) | N = 751; Harems = 61; Sites = 8 |
| **6** | LMER | Gaussian | Long tail | log(0.3 +Interaction) ~ Scenario * days + (1\|Site/ID) | N = 70 (Alive: 34; Dead: 38); Cleaners = 23 |
| **7** | GLM | Binomial (logit) | Bounded proportions | Prop ~ Strategy*LifeHistory, weights | N = 48; Sites = 8; Strategy = 3; LifeHistory = 2 |
| **8** | GLM | Binomial (logit) | Bounded proportions | Prop~ Strategy * Density, weights | N = 24; Sites = 8; Strategy = 3 |
| **9** | LM | Gaussian | - | Size ~ Migration * Scenario | N = 42 |
| **10** | LM | Gaussian | - | Size ~ Density * Scenario | N = 42 |

### Supplement 4: Details for Model 1

#### Table S4.1: Wilcoxon signed rank test with continuity correction

| Model | Sex | V | P value |
|---|---|---|---|
| M1 | All | 5291.5 | 4.75e-06 |
| M1.1 | Males | 730.5 | 0.038 |
| M1.2 | Females | 2109.5 | 2.845e-05 |
| | Alternative Hypothesis: true location shift is not equal to 0 | | |

### Supplement 5: Details for Model 2

#### Figure S5: Diagnostic Plots

The figure below shows the assumption checks for Model 2 (LMER):

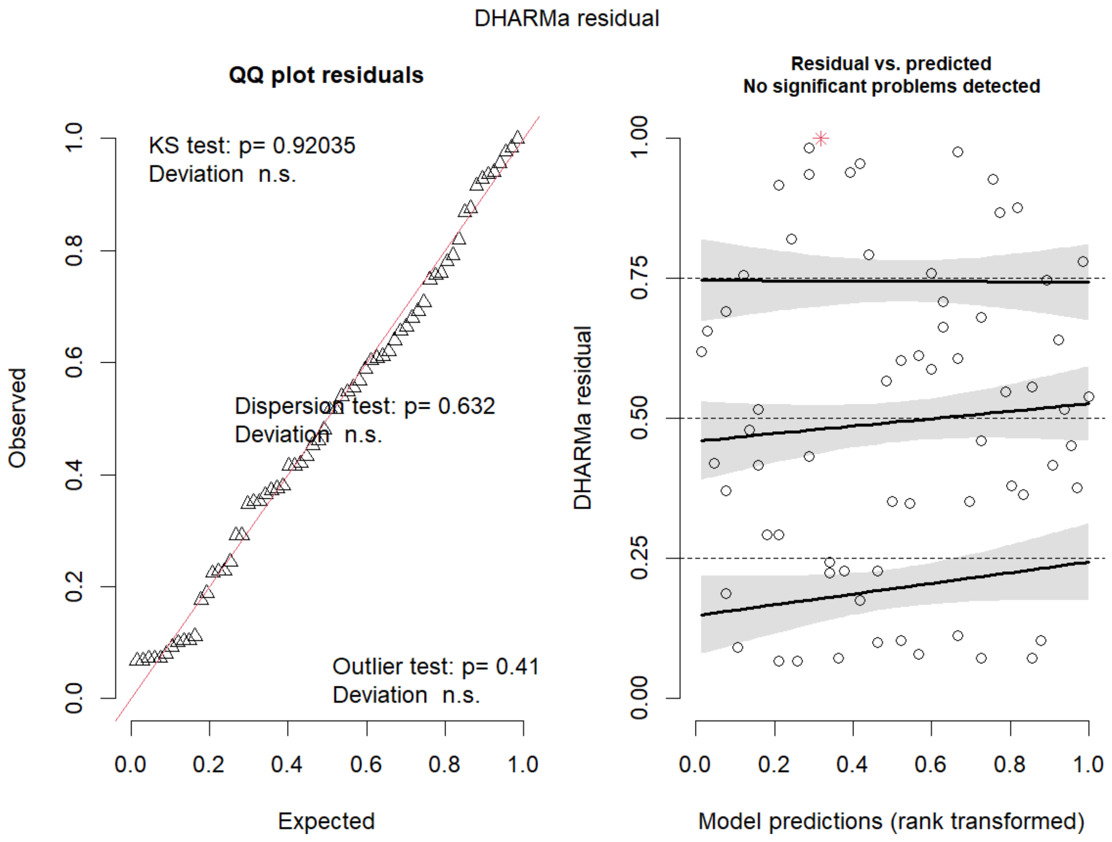

**Table S5.1: Analysis of Deviance – Type II Wald Chi-square Tests**

| Term | Chisq | Df | Pr(>Chisq) |
|---|---|---|---|
| Sex | 4.9934 | 1 | 0.025442 * |
| Scenario | 0.3043 | 1 | 0.5811872 |
| Initial-Size | 12.2010 | 1 | 0.0004777*** |
| Sex : Scenario | 0.0781 | 1 | 0.7798972 |
| Sex : Initial-Size | 1.2521 | 1 | 0.2631450 |

**Table S5.2: Emmeans Contrasts**

| Initial Size | Ratio | SE | Df | T ratio | P value |
|---|---|---|---|---|---|
| | | Contrast Female / Male | | | |
| 60 | 0.903 | 0.456 | 1 | -0.201 | 0.8417 |
| 61 | 0.932 | 0.447 | 1 | -0.146 | 0.8847 |
| 62 | 0.962 | 0.436 | 1 | -0.084 | 0.9332 |
| 63 | 0.993 | 0.425 | 1 | -0.015 | 0.9878 |
| 64 | 1.025 | 0.413 | 1 | 0.062 | 0.9506 |
| 65 | 1.059 | 0.400 | 1 | 0.150 | 0.8814 |
| 66 | 1.093 | 0.387 | 1 | 0.250 | 0.8040 |
| 67 | 1.128 | 0.373 | 1 | 0.364 | 0.7184 |
| 68 | 1.164 | 0.359 | 1 | 0.493 | 0.6252 |
| 69 | 1.202 | 0.345 | 1 | 0.641 | 0.5261 |
| 70 | 1.241 | 0.331 | 1 | 0.809 | 0.4247 |
| 71 | 1.281 | 0.317 | 1 | 0.998 | 0.3261 |
| 72 | 1.322 | 0.306 | 1 | 1.207 | 0.2367 |
| 73 | 1.364 | 0.296 | 1 | 1.432 | 0.1622 |
| 74 | 1.408 | 0.290 | 1 | 1.664 | 0.1062 |
| 75 | 1.454 | 0.288 | 1 | 1.889 | 0.0682 |
| 76 | 1.501 | 0.292 | 1 | 2.090 | 0.0449 |
| 77 | 1.549 | 0.301 | 1 | 2.250 | 0.0316 |
| 78 | 1.599 | 0.318 | 1 | 2.361 | 0.0247 |
| 79 | 1.651 | 0.342 | 1 | 2.421 | 0.0215 |
| 80 | 1.704 | 0.372 | 1 | 2.438 | 0.0207 |

Re75sults averaged over the levels of Scenario, Degrees of freedom method: Kenward-Roger. Results are given on the log scale (not the response one).

### Supplement 6: Details for Model 3

#### Figure S6: Diagnostic Plots

The figure below shows the assumption checks for Model 3 (LMER):

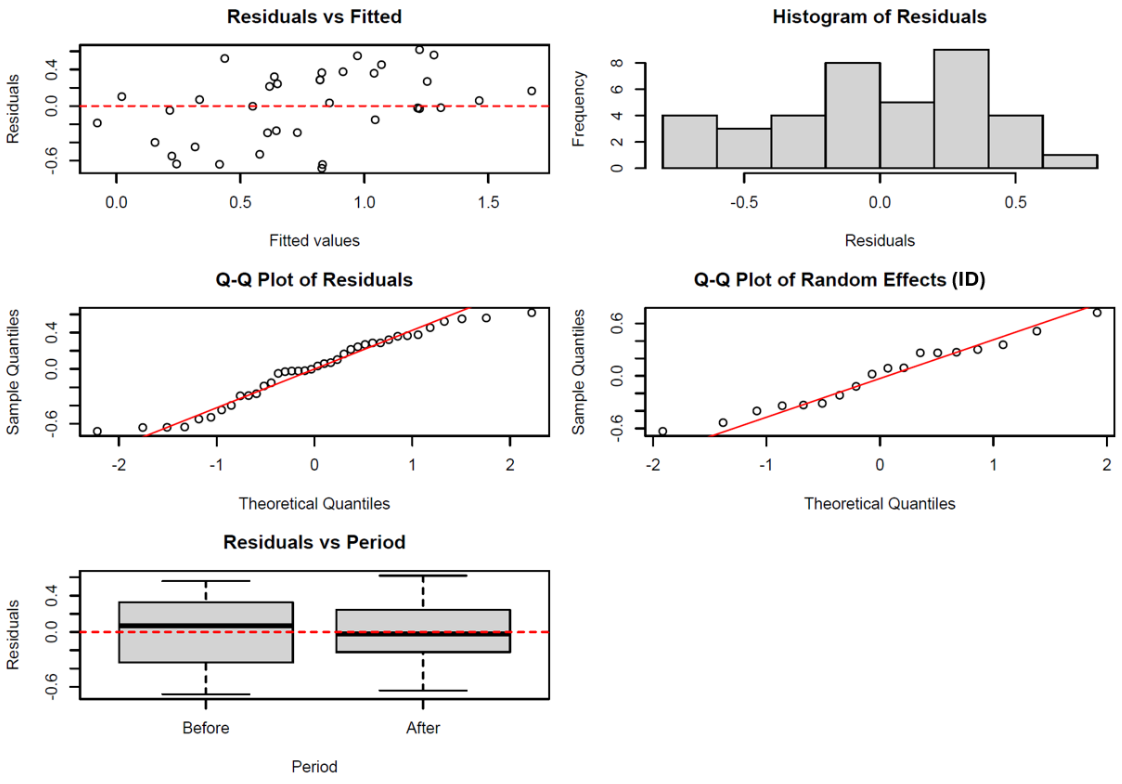

#### Table S6.1: Analysis of Deviance – Type II Wald Chi-square Tests

| Term | Chisq | Df | Pr(>Chisq) |
|---|---|---|---|
| Period | 6.8265 | 1 | 0.00891 ** |

#### Table S6.2: Emmeans contrasts

| Contrast | Estimate | SE | df | t.ratio | p-value |
|---|---|---|---|---|---|
| Period | 0.394 | 0.151 | 19 | 2.613 | 0.0171 |

### Supplement 7: Details for Model 4

#### Figure S7: Diagnostic Plots

The figure below shows the assumption checks for Model 4 (GLM with Poisson family):

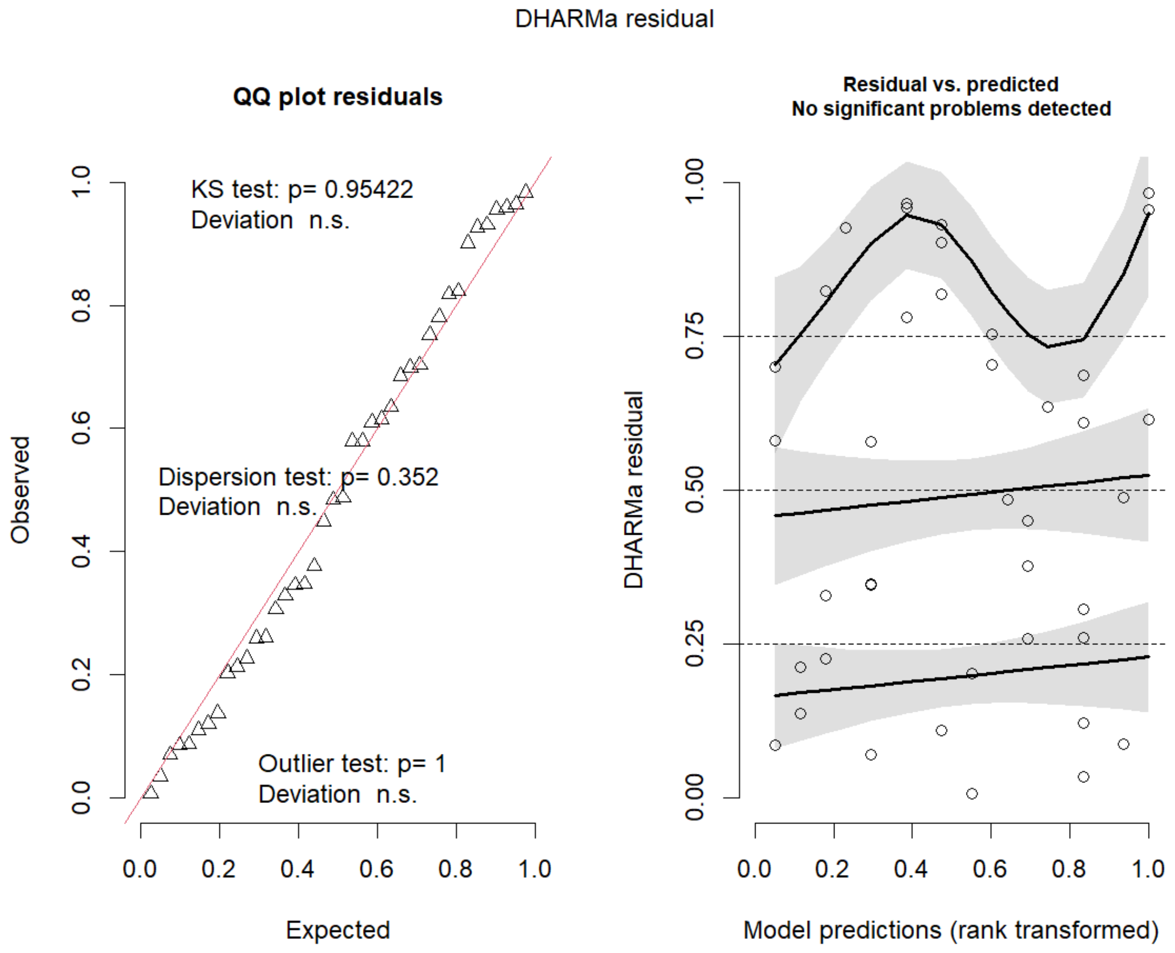

#### Table S7: Analysis of Deviance – Type II Wald Chi-square Tests

| Term | LR Chisq | Df | Pr(>Chisq) |
|---|---|---|---|
| System | 25.3508 | 1 | 4.78e-07 *** |
| Density | 1.1335 | 1 | 0.28702 . |
| System : Density | 3.2571 | 1 | 0.07112 |

### Supplement 8: Details for Model 5

#### Figure S8: Diagnostic Plots

The figure below shows the assumption checks for Model 1 (GLM with Poisson family):

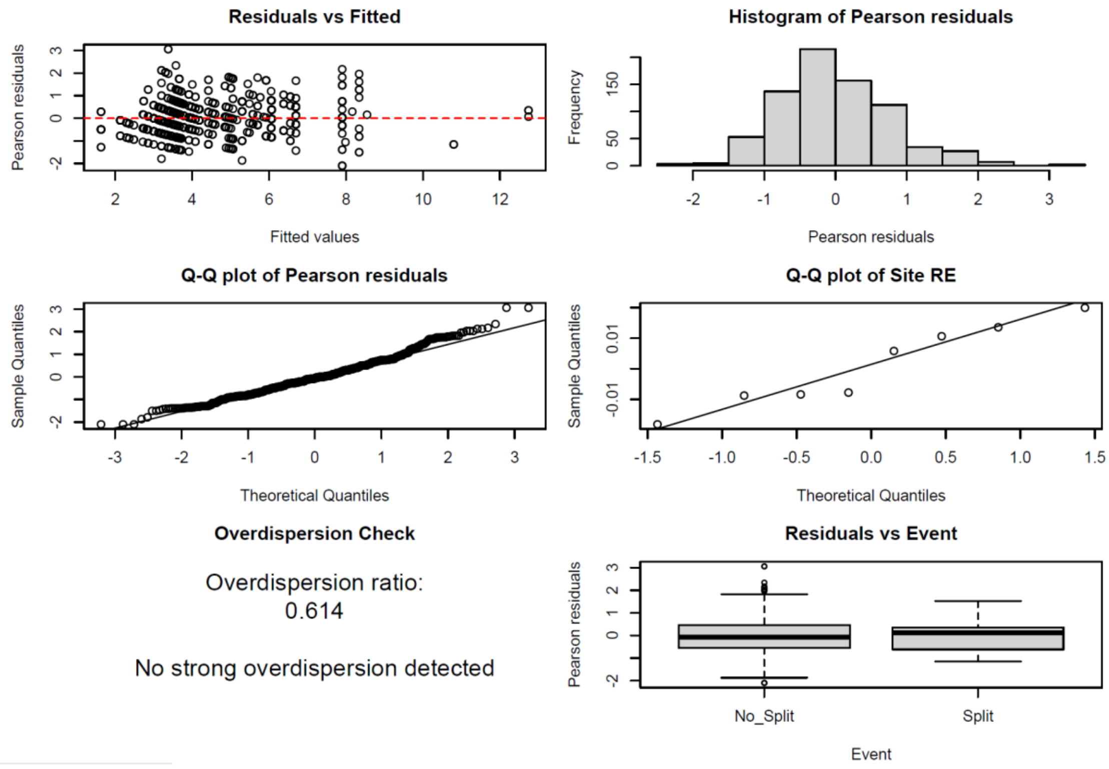

#### Table S8: Analysis of Deviance – Type II Wald Chi-square Tests

| Term | LR Chisq | Df | Pr(>Chisq) |
|---|---|---|---|
| Splitting | 23.996 | 1 | 9.654e-07 *** |

### Supplement 9: Details for Model 6

#### Figure S9: Diagnostic Plots

The figure below shows the assumption checks for Model 6 (LMER):

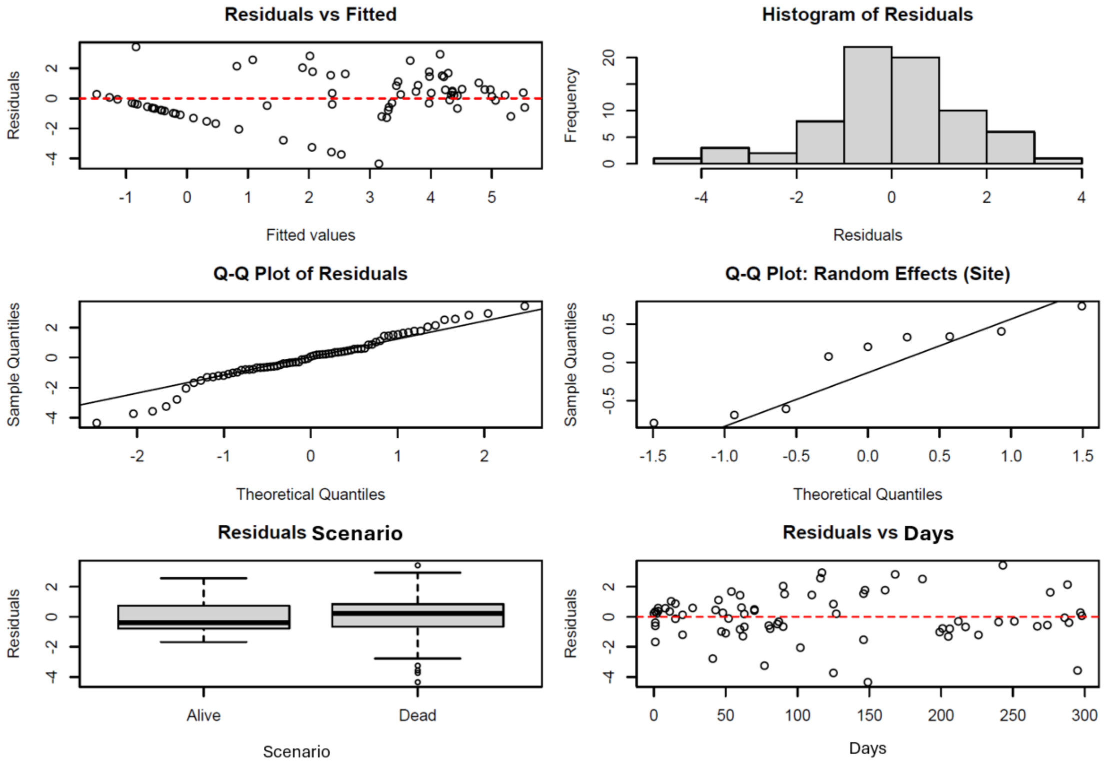

#### Table S9.1: Analysis of Deviance – Type II Wald Chi-square Tests

| Term | Chisq | Df | Pr(>Chisq) |
|---|---|---|---|
| Scenario | 7.9220 | 1 | 0.004884 ** |
| Days | 15.2425 | 1 | 9.455e-05 *** |
| Scenario : Days | 0.0143 | 1 | 0.904960 |

#### Table S9.2: Emmeans contrasts

| Scenario | emmean | Back transformed emmean | SE | df |
|---|---|---|---|---|
| Early | 1.088395 | 2.042953 | 0.7143962 | 10.66 |
| Standard | 3.156595 | 23.190463 | 0.6916696 | 9.93 |

### Supplement 10: Details for Model 7

#### Figure S10: Diagnostic Plots

The figure below shows the assumption checks for Model 7 (GLM with binomial family):

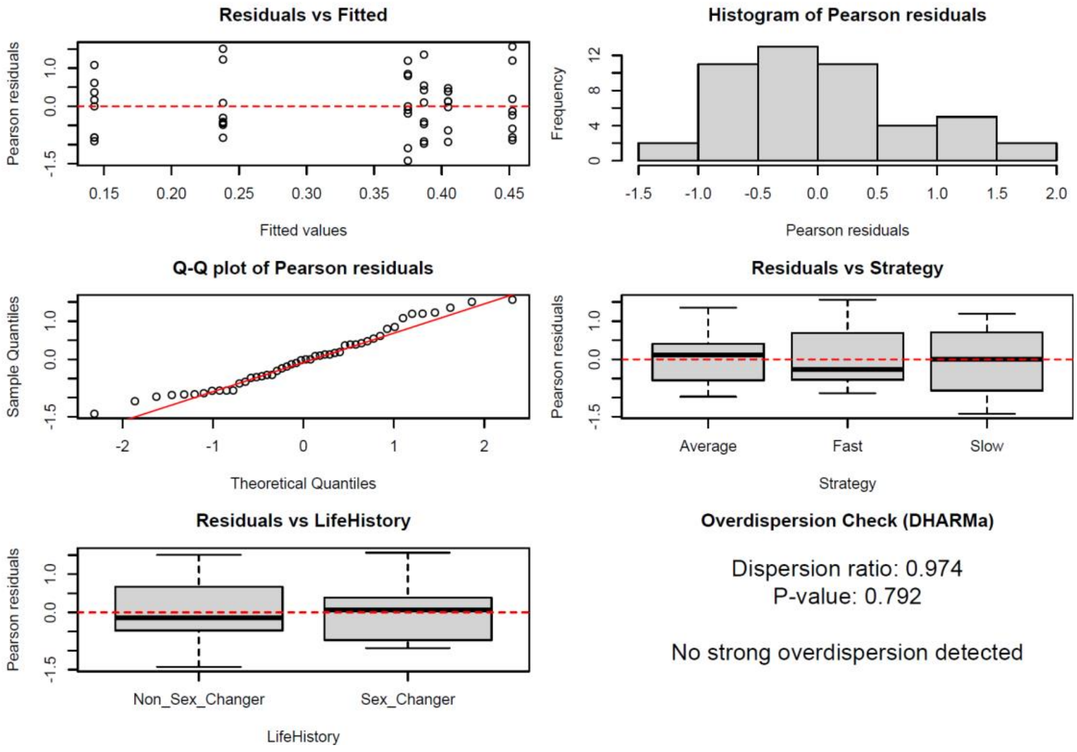

#### Table S10.1: Analysis of Deviance – Type II Wald Chi-square Tests

| Term | LR Chisq | Df | Pr(>Chisq) |
|---|---|---|---|
| Strategy | 14.591 | 2 | 0.0006786 *** |
| LifeHistory | 0 | 1 | 0.9999998 |
| Strategy : LifeHistory | 18.171 | 2 | 0.0001133 *** |

#### Table S10.2: Emmeans Contrasts

| | Estimate | SE | Df | Z ratio | P value |
|---|---|---|---|---|---|
| Average | -0.0747 | 0.334 | Inf | -0.224 | 0.8229 |
| Fast | -0.9721 | 0.335 | Inf | -2.898 | 0.0038 |
| Slow | 1.2809 | 0.455 | inf | 2.814 | 0.0049 |

Results are given on the log odd ratio scale (not response one)

### Supplement 11: Details for Model 8

#### Figure S11: Diagnostic Plots

The figure below shows the assumption checks for Model 8 (GLM with binomial family):

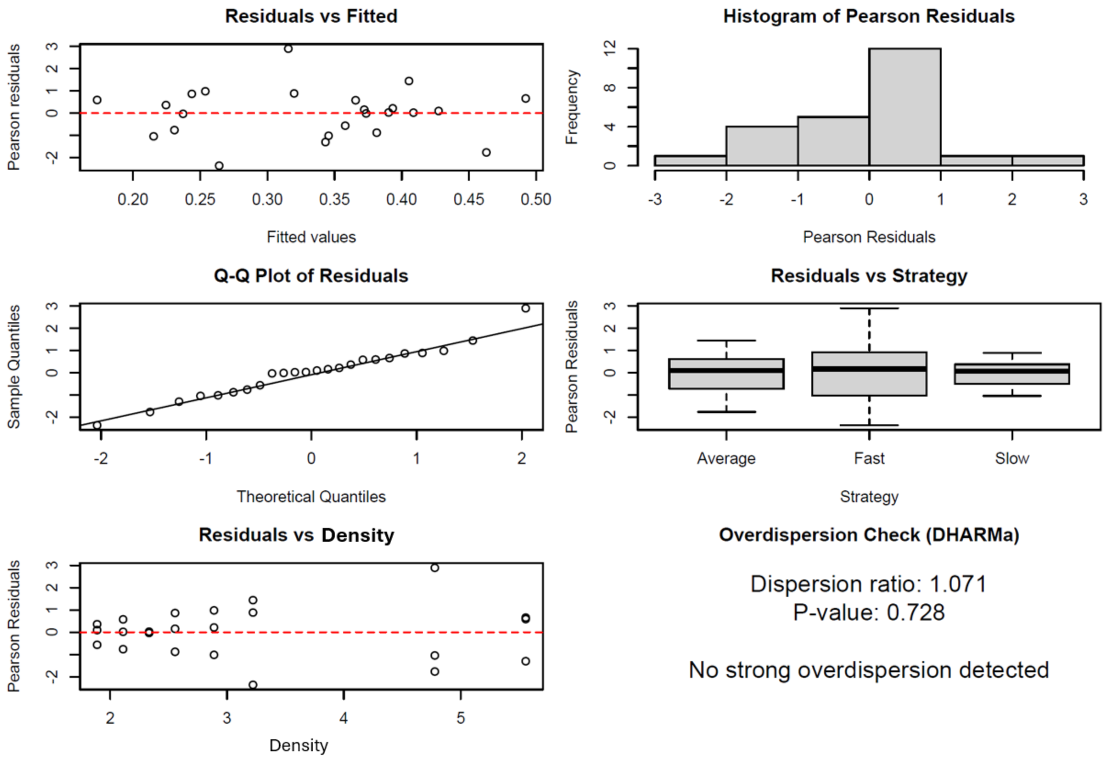

#### Table S11.1: Analysis of Deviance – Type II Wald Chi-square Tests

| Term | LR Chisq | Df | Pr(>Chisq) |
|---|---|---|---|
| Strategy | 16.760 | 2 | 0.0002294 *** |
| Density | 0 | 1 | 0.999999 |
| Strategy : Density | 19.787 | 2 | 0.05e-05 *** |

#### Table S11.2: Emmtrends

| | Density trend | SE | Df | Asymp LCL | Asymp UCL |
|---|---|---|---|---|---|
| Average | 0.151 | 0.123 | Inf | -0.0131 | 0.315 |
| Fast | 0.161 | 0.0907 | Inf | -0.0168 | 0.339 |
| Slow | -0.346 | 0.996 | inf | -0.5417 | -0.151 |

### Supplement 12: Details for Model 9

#### Figure S12: Diagnostic Plots

The figure below shows the assumption checks for Model 9 (LM):

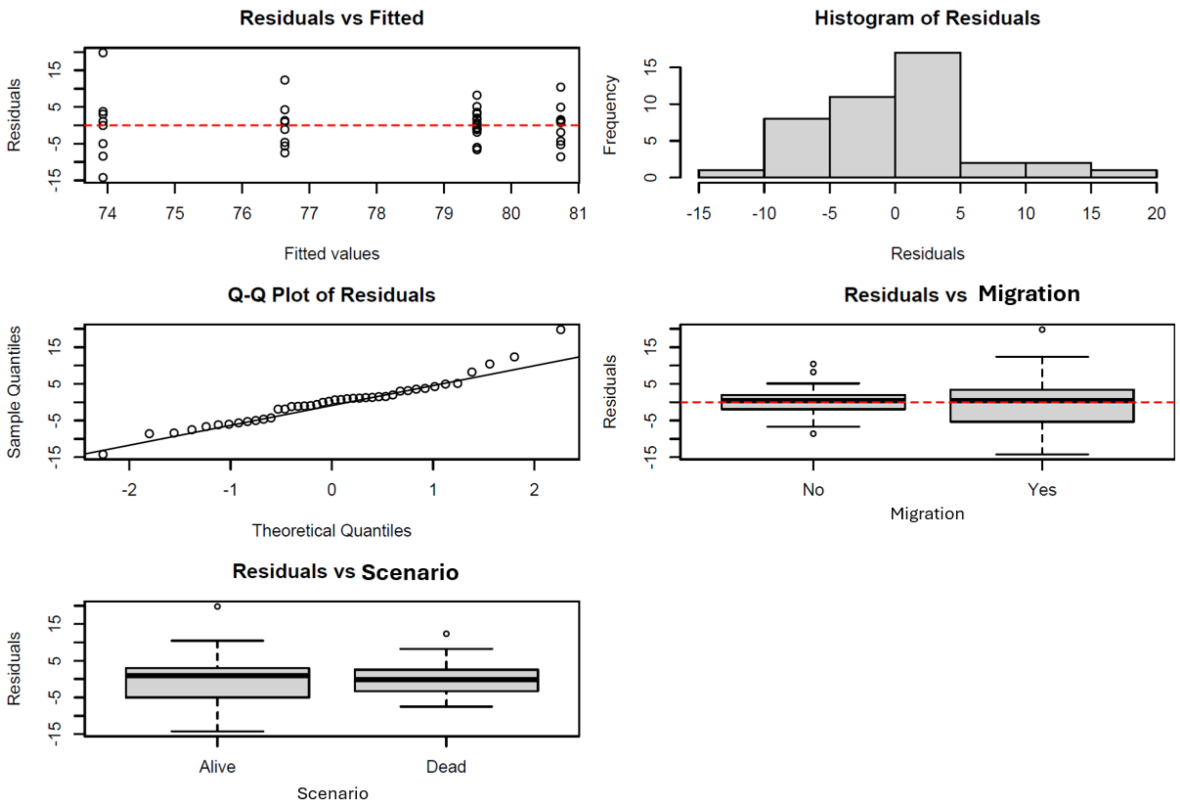

#### Table S12: Analysis of Deviance Table (Type II Tests)

| Term | Sum Sq | Df | F value | Pr(>F) |
|---|---|---|---|---|
| Migration | 211.60 | 1 | 5.3264 | 0.02654 * |
| Scenario | 0.98 | 1 | 0.0246 | 0.87632 |
| Migration : Scenario | 37.81 | 1 | 0.9517 | 0.33545 |
| Residuals | 1509.58 | 38 | | |

### Supplement 13: Details for Model 10

#### Figure S13: Diagnostic Plots

The figure below shows the assumption checks for Model 10 (LM ):

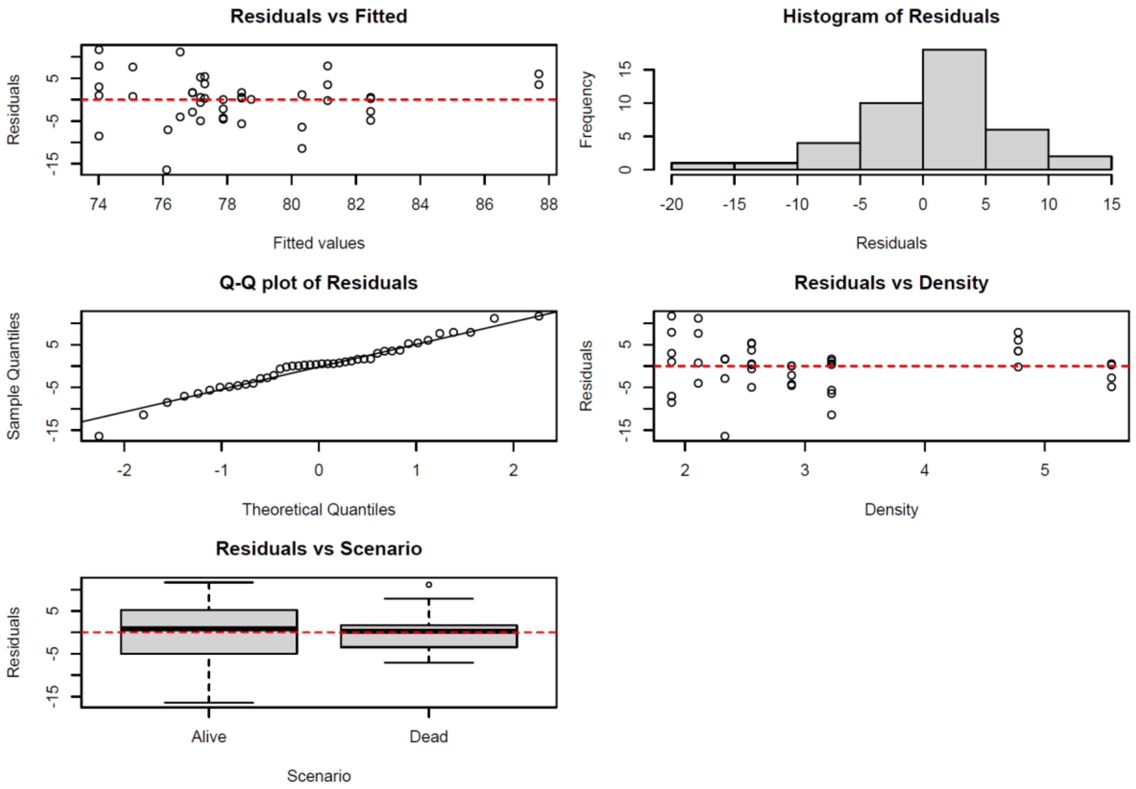

#### Table S13: Analysis of Deviance Table (Type II Tests)

| Term | Sum Sq | Df | F value | Pr(>F) |
|---|---|---|---|---|
| Density | 323.95 | 1 | 9.3530 | 0.004065 ** |
| Scenario | 6.71 | 1 | 0.1937 | 0.662349 |
| Density : Scenario | 90.46 | 1 | 2.6117 | 0.114351 |
| Residuals | 1316.15 | 38 | | |

### Supplement 14: Details on sex changers in the alive and dead scenario

#### Table S14.1.1: Information on the sex changing females in the alive scenario

| Sex Changer | Social System | Migration to periphery | Codominant females | Map appendix |
|---|---|---|---|---|
| ***Alive Scenario*** | | | | |
| CGRFYY | Branching | No | 2 | S14.1a |
| HLMRR | Linear | Yes | NA | S14.1b |
| LUAnninuccia | Linear | Yes | 0 | S14.1c |
| LUWhiteWoman | Branching | No | 2 | S14.1d |
| MLApprentist2 | Linear | No | 0 | S14.1e |
| MLLFYR | Linear | Yes | 0 | S14.1f |
| MLLMP | Branching | No | 4 | S14.1g |
| MLNewRFWW | Branching | No | 4 | S14.1g |
| MLRMYY | Branching | No | 4 | S14.1g |
| MRLMWW | Branching | Yes | 2 | S14.1h |
| MRPuntino | Branching | No | 3 | S14.1i |
| MRRMGR | Branching | No | 3 | S14.1i |
| MRRMRY | Branching | No | 3 | S14.1j |
| ONewBBFemale | Branching | No | 3 | S14.1k |
| ORFPY | Branching | Yes | 3 | S14.1k |
| OWhiteNinja | Branching | No | 2 | S14.1l |
| WLMGG | Branching | No | 2 | S14.1m |
| WRFYY | Branching | No | 3 | S14.1m |

#### Table S14.1.2: Maps of Sex changing females and their male in the alive scenario

The following table shows the territories of the sex changer (in pink), her male (in blue) and potential co-dominant females (in yellow, purple), before (left side) and after (right side) the sex change event.

<table>
<tr><th>BEFORE SEX CHANGE</th><th>AFTER SEX CHANGE</th></tr>
<tr><td colspan="2">S14.1a : CGRFYY</td></tr>
<tr><td>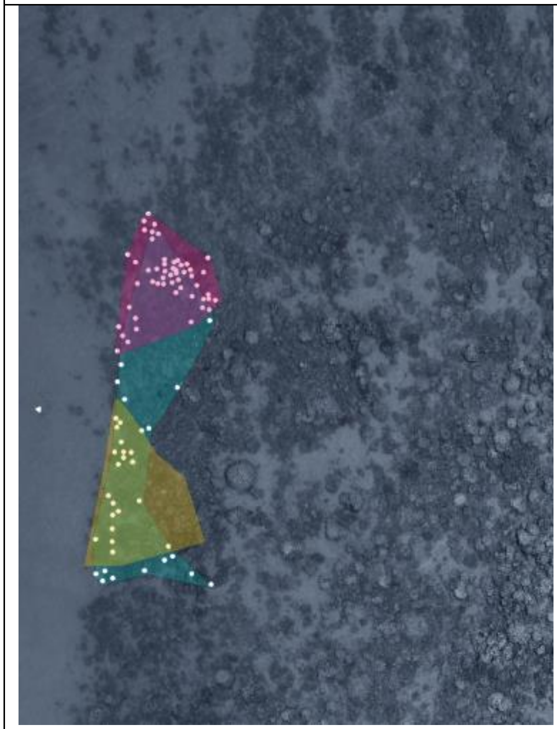</td><td>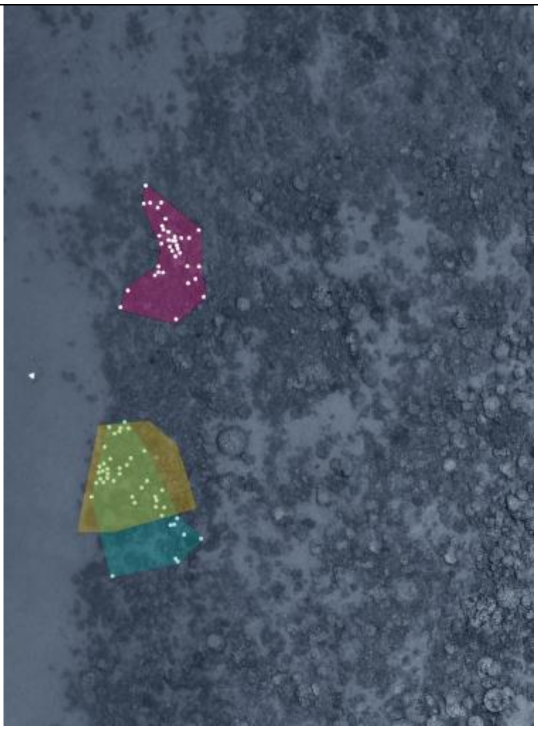</td></tr>
<tr><td colspan="2">S14.1d : LUWhiteWoman</td></tr>
<tr><td>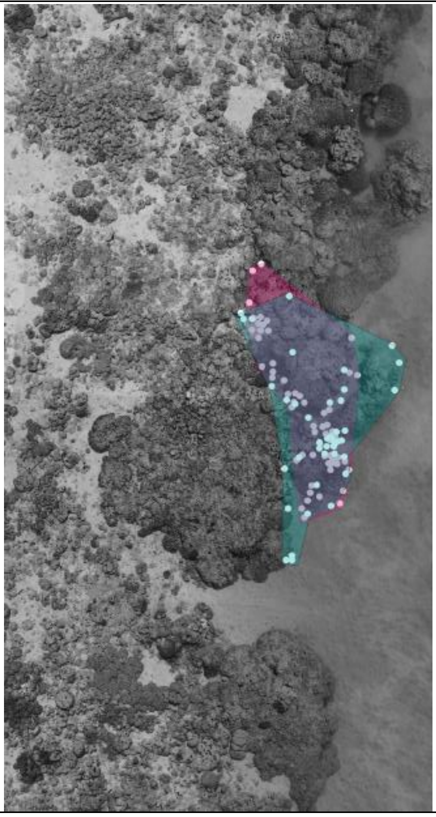</td><td>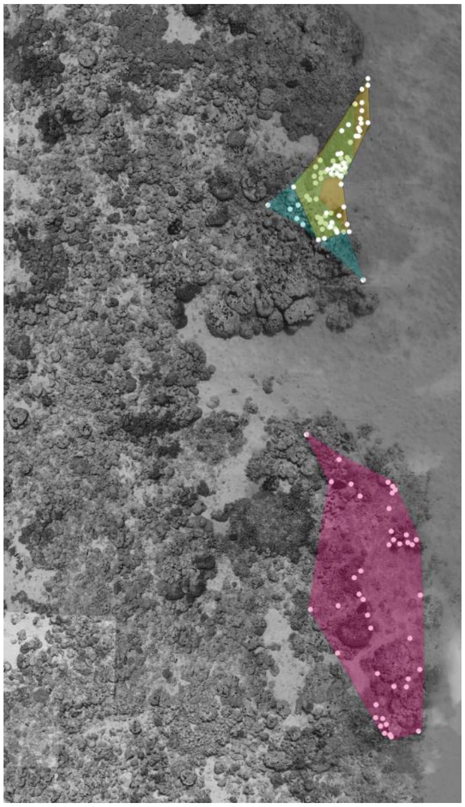</td></tr>
</table>

| **S14.1k : OnewBBFemale, ORFPY** | |
|---|---|
| 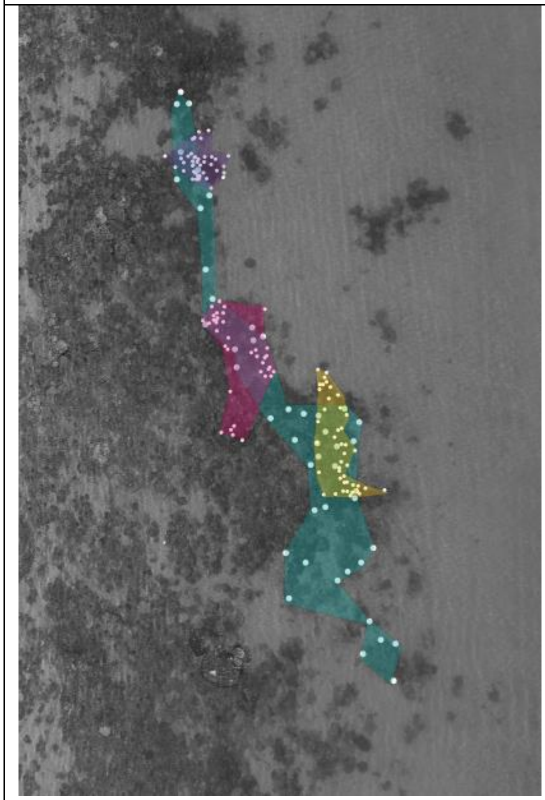 | 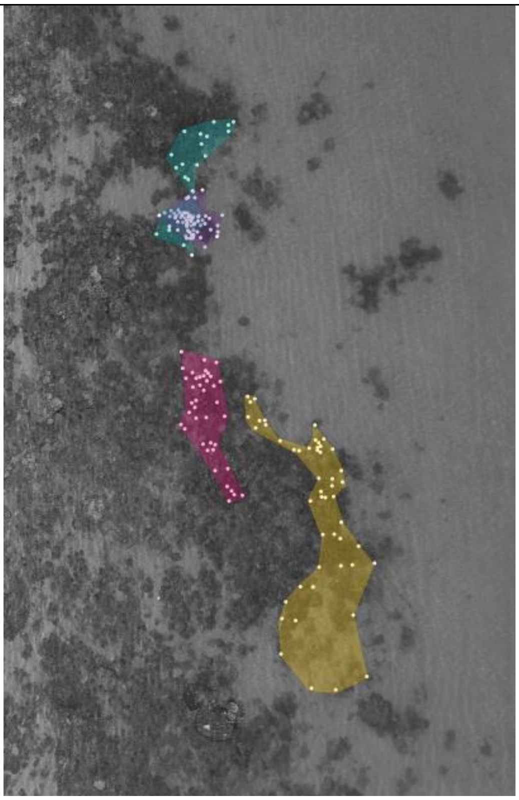 |
| **S14.1l : OWhiteNinja** | |
| 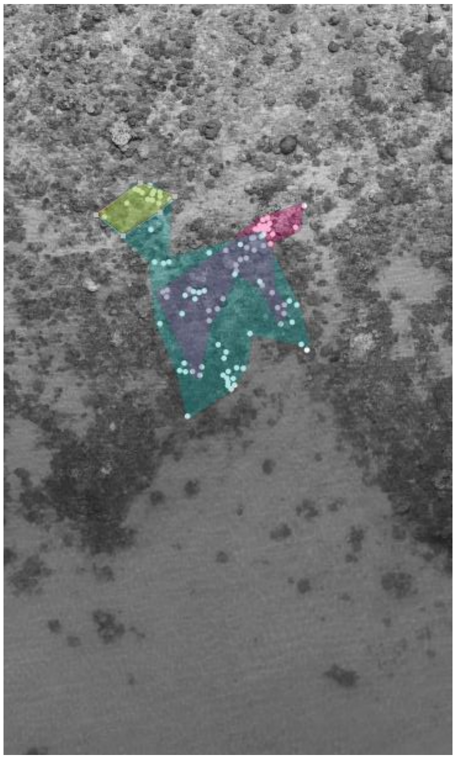 | 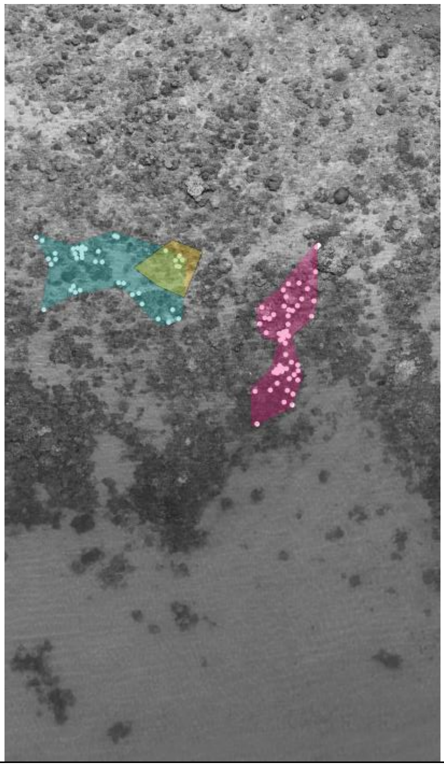 |

S14.1g:  MLRFWW, MLLMP; MLRMYY

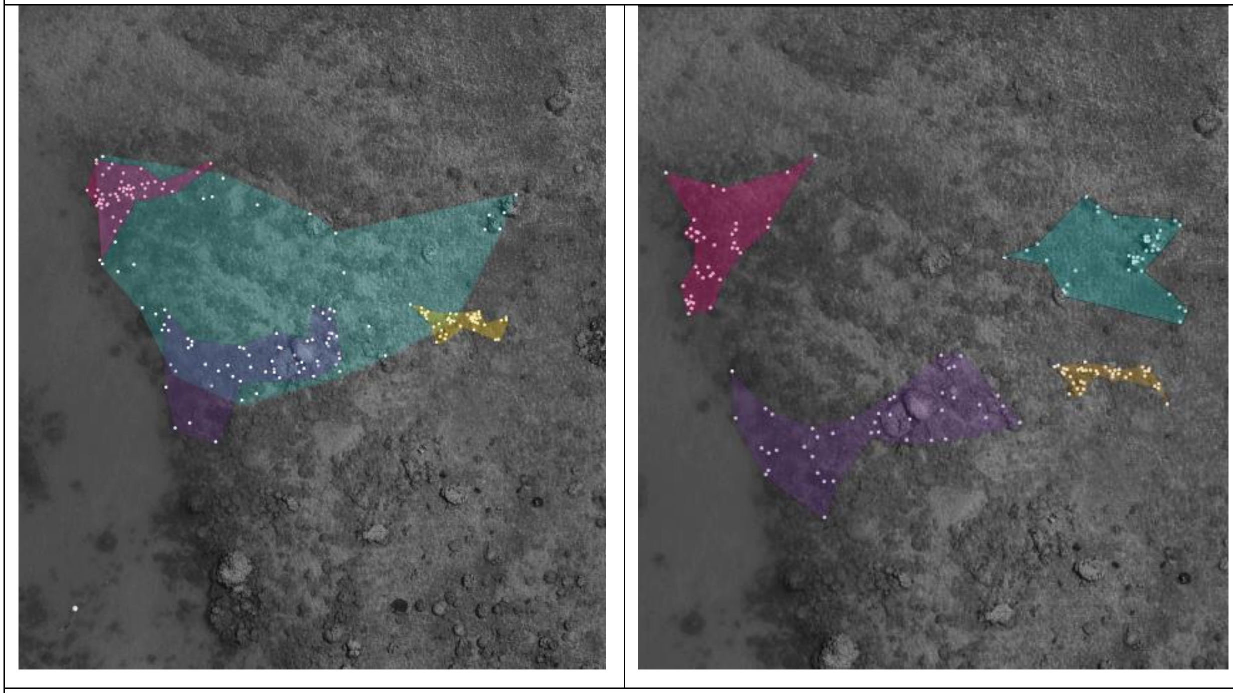

S14.1j:  MRRMRY

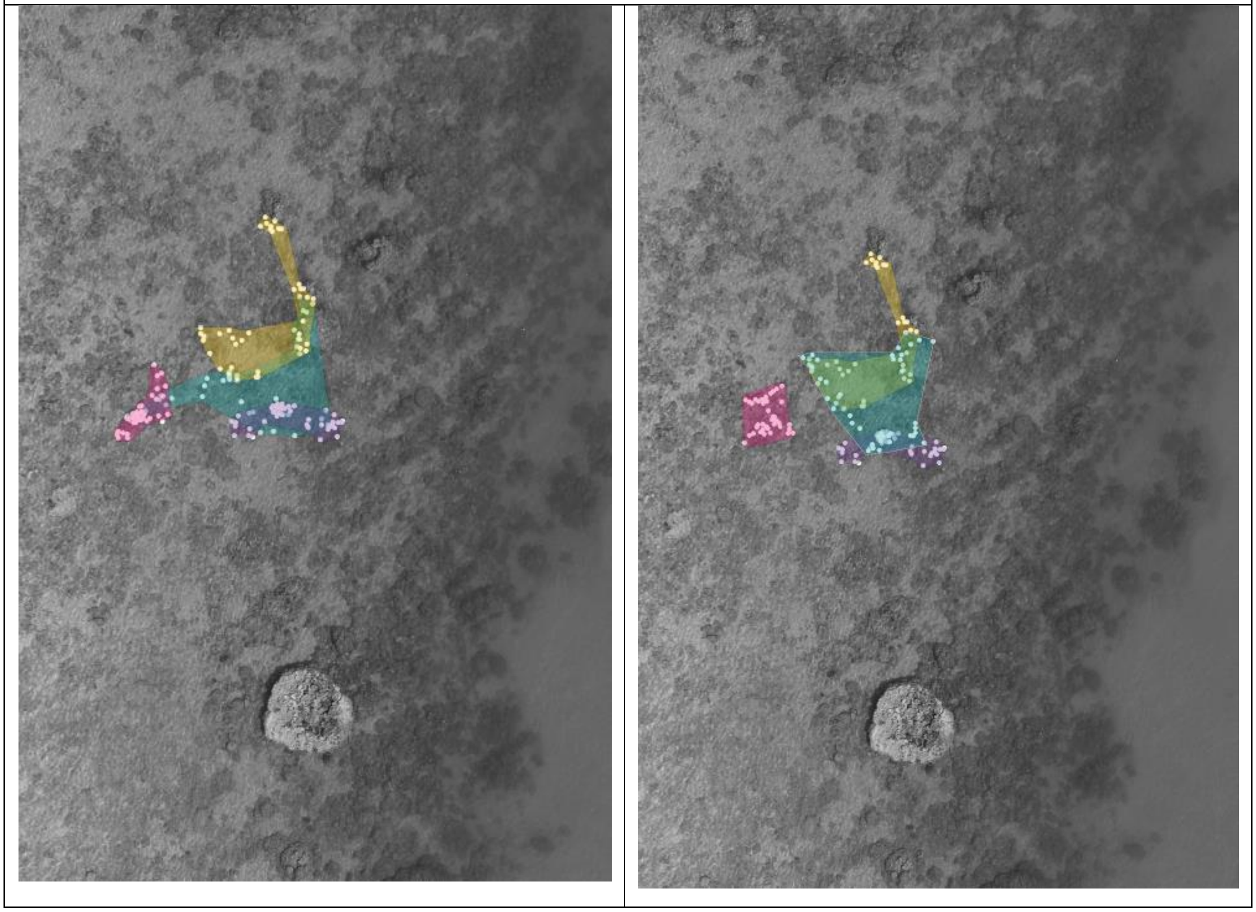

| S14.1m: WRFYY, WLMGG | |
|---|---|
| 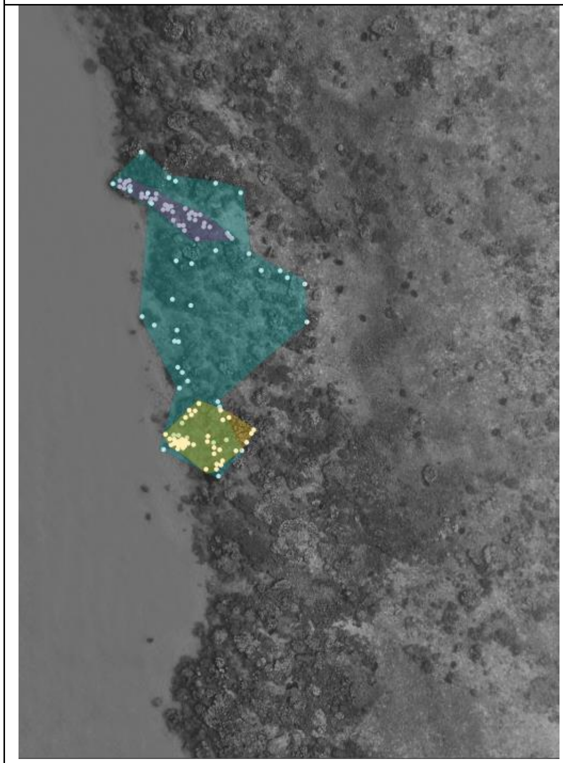 | 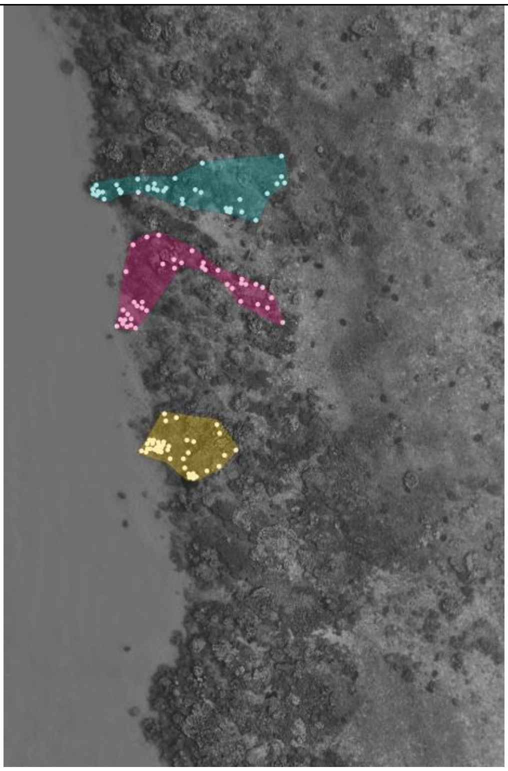 |
| **S14.1c: LUAnninuccia** | |
| 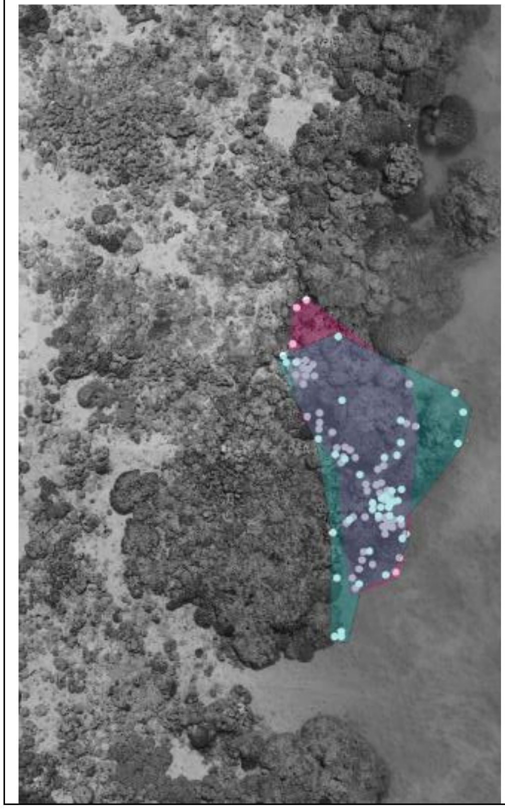 | 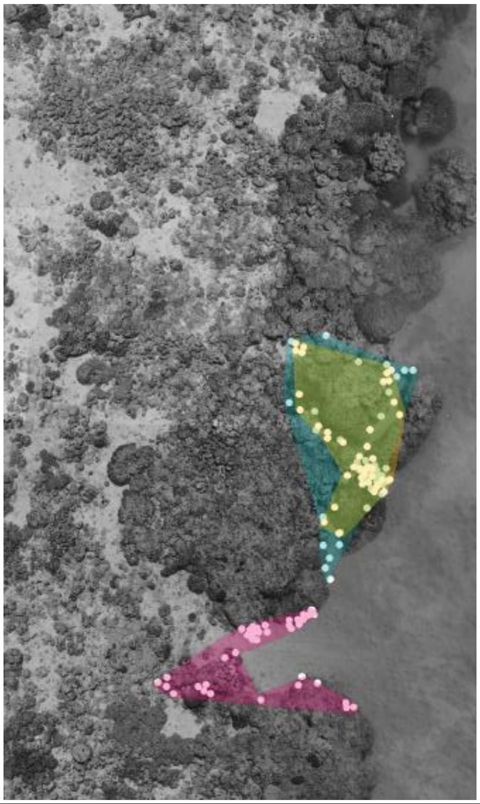 |

**S14.1i: MRPuntino, MRRMGR**

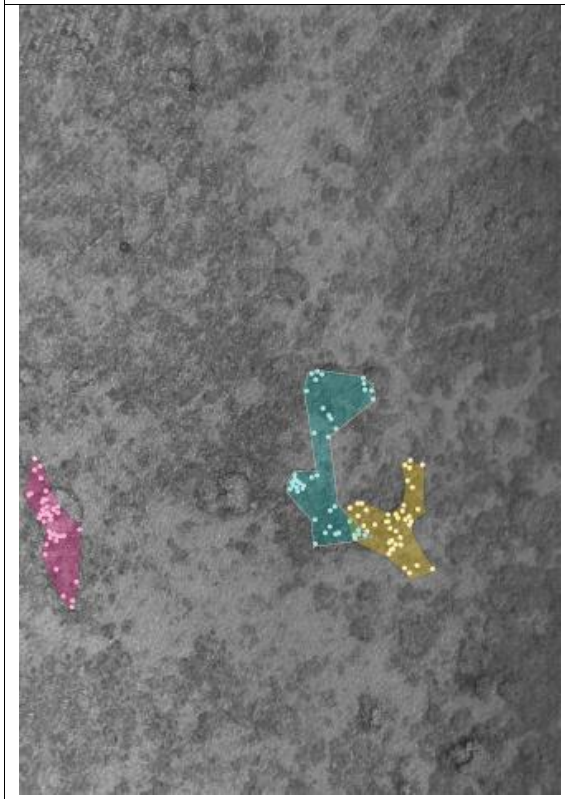

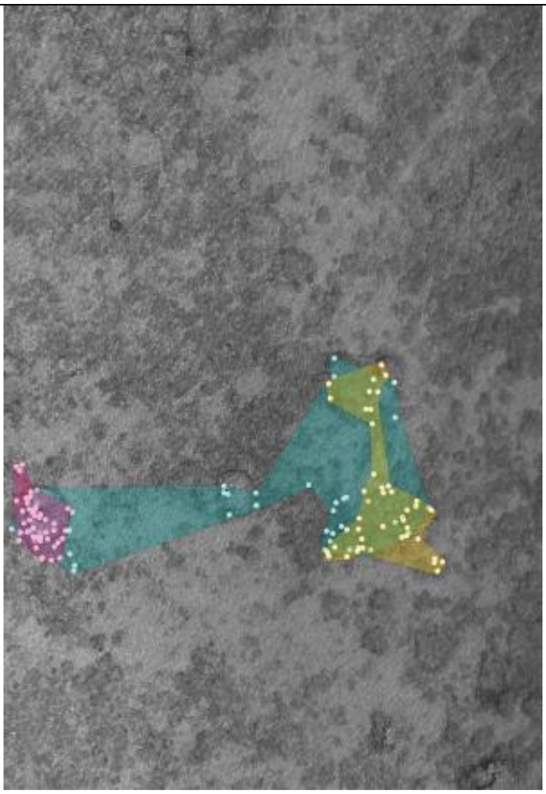

**S14.1h: MRLMWW**

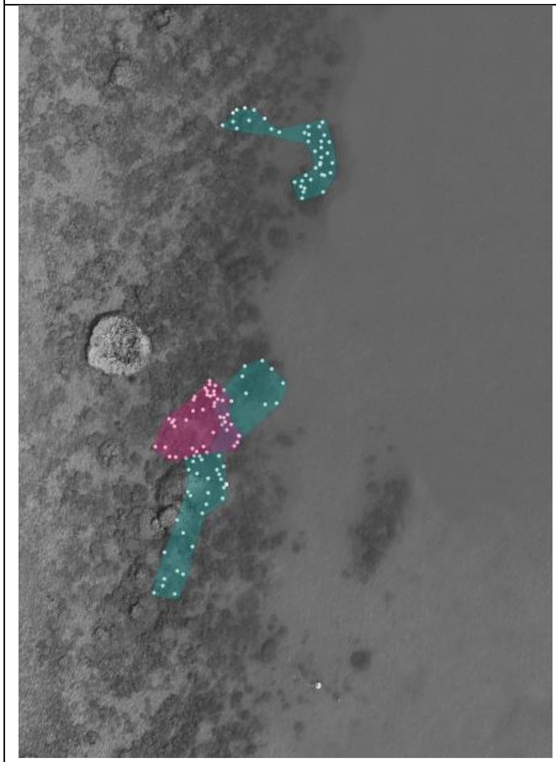

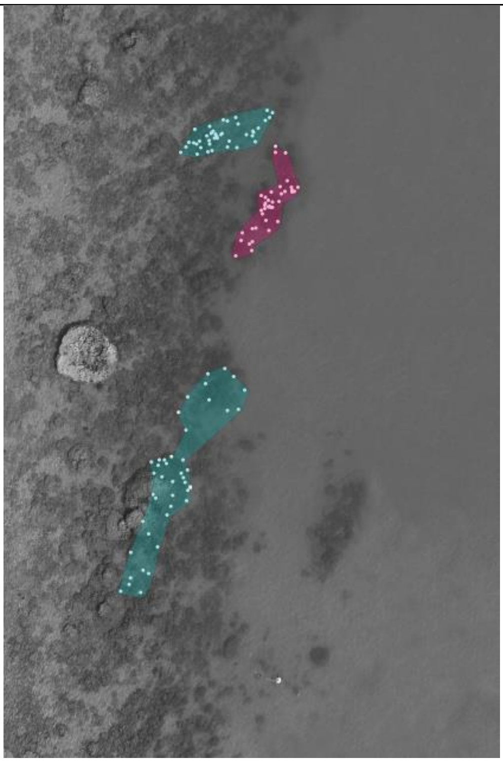

**S14.1f: MLLFYR**

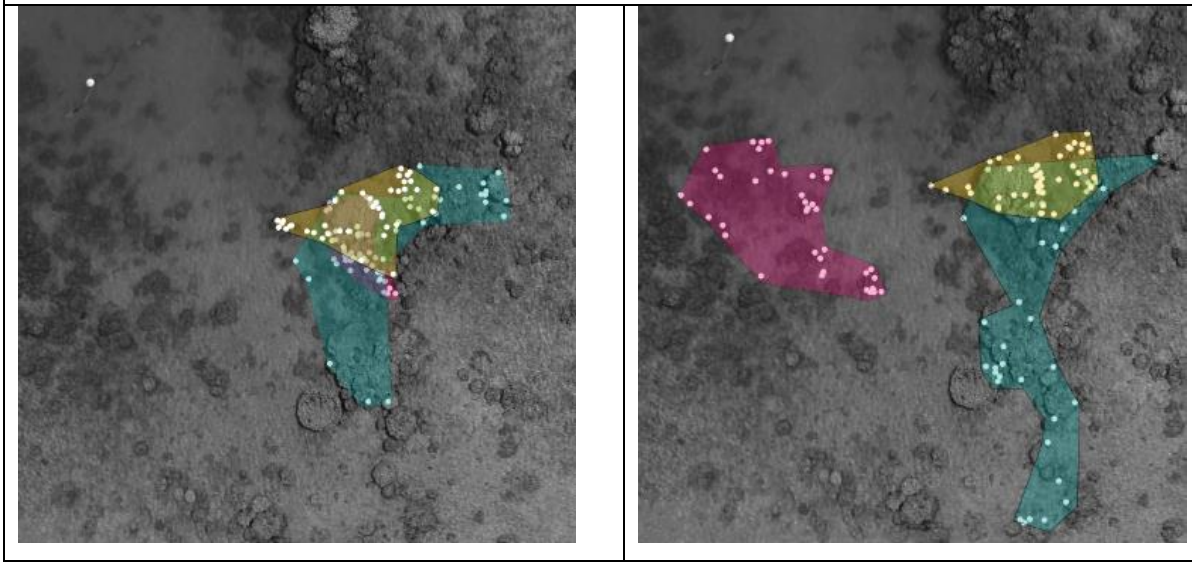

### Table S14.2.1: Information on the sex changing females in the dead scenario

| Sex Changer | Social System | Migration to periphery | Codominant females | Map appendix |
|---|---|---|---|---|
| ***Dead Scenario*** | | | | |
| CGLFGG | Linear | No | 0 | S14.2a |
| CGLFPY | Linear | No | 0 | S14.2b |
| CGMinnie | Linear | No | 0 | S14.2c |
| CGNewLFPP | Linear | No | 0 | S14.2d |
| HLFWW | Linear | No | 0 | S14.2e |
| HRMYY | Linear | No | 0 | S14.2f |
| Hyelcor | Linear | No | 0 | S14.2g |
| LUAnnFriend | Linear | No | 0 | S14.2h |
| LUBS2 | Linear | No | 0 | S14.2i |
| LUWhitie | Linear | No | 0 | S14.2j |
| MBB2Alpha | Linear | No | 0 | S14.2k |
| MBIslGirl | Branching | No | 2 | S14.2l |
| MBLMPGFriend | Linear | No | 0 | S14.2m |
| MBWierdo | Branching | No | 2 | S14.2l |
| MLApprentist | Linear | No | 0 | S14.2n |
| MRAlphaRR | Branching | No | 2 | S14.2o |
| MRLMRY | Linear | No | 0 | S14.2p |
| MRNewLFWW | Branching | No | 2 | S14.2o |
| OLMPY | Linear | No | 0 | S14.2q |
| ORFGGMP | Linear | No | 0 | S14.2r |
| ORFPP | Linear | No | 0 | S14.2s |
| ORFWP | Linear | No | 0 | S14.2t |
| WLFPY | Linear | No | 0 | S14.2u |

### Table S14.2.2: Maps of Sex changing females and their male in the dead scenario

The following table shows the territories of the sex changer (in pink), her male (in blue) and potential co-dominant females (in yellow, purple), before (left side) and after (right side) the sex change event.

<table>
<tr><th>BEFORE SEX CHANGE</th><th>AFTER SEX CHANGE</th></tr>
<tr><td colspan="2">S14.2b : CGLFPY</td></tr>
<tr><td>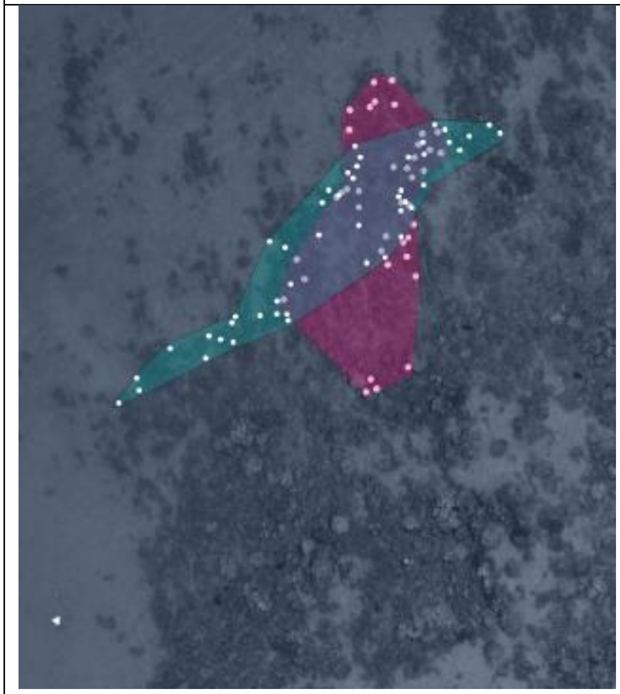</td><td>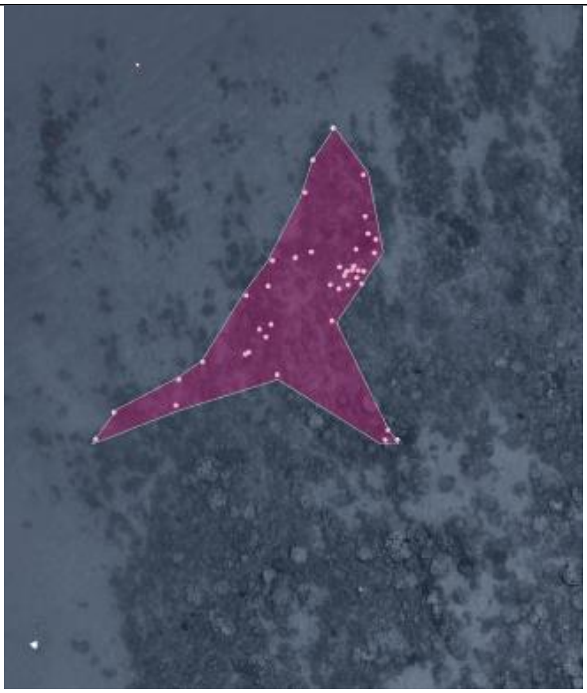</td></tr>
<tr><td colspan="2">S14.2c: CGMinnie</td></tr>
<tr><td>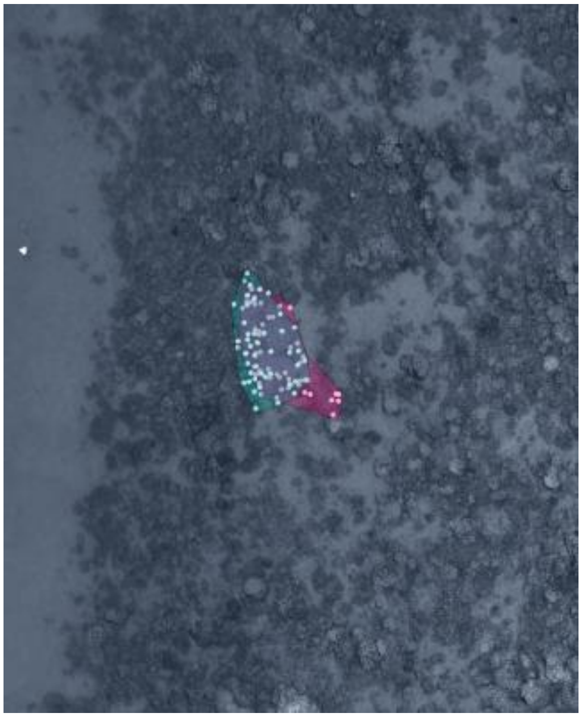</td><td>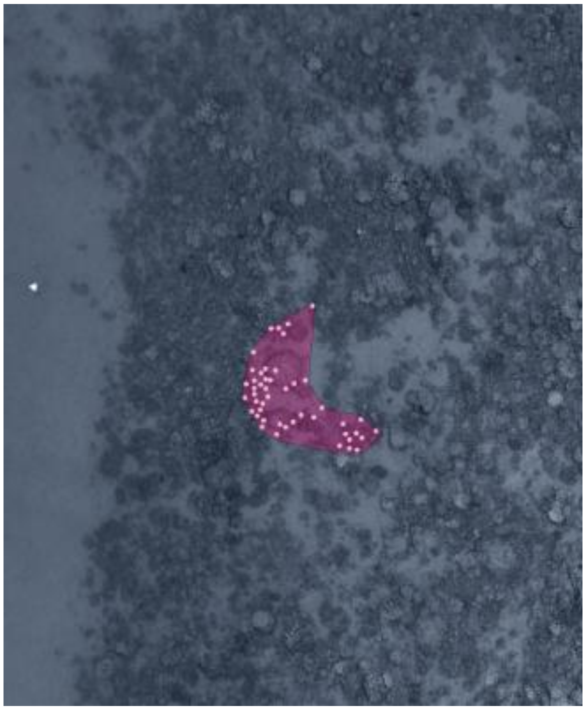</td></tr>
</table>

| | |
|---|---|
| **S14.2d: CGNewLFPP** | |
| 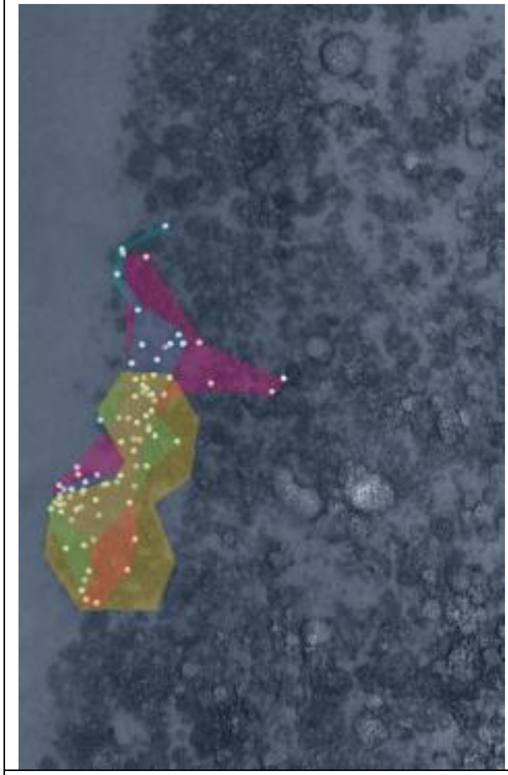 | 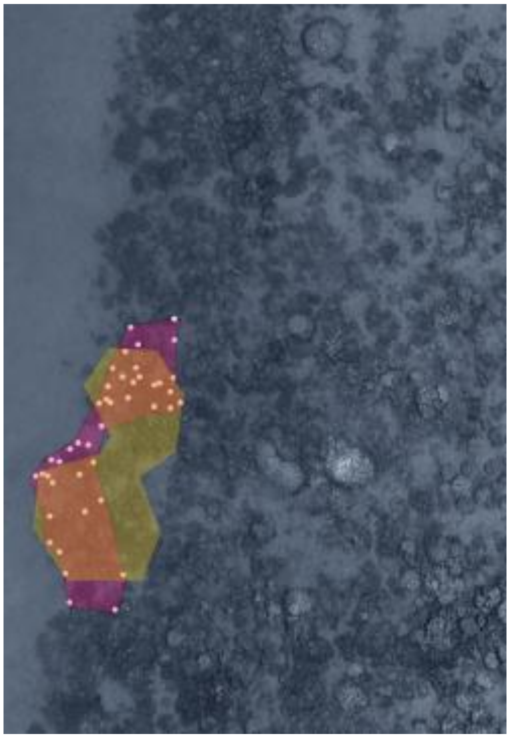 |
| **S14.2j: LUWhitie** | |
| 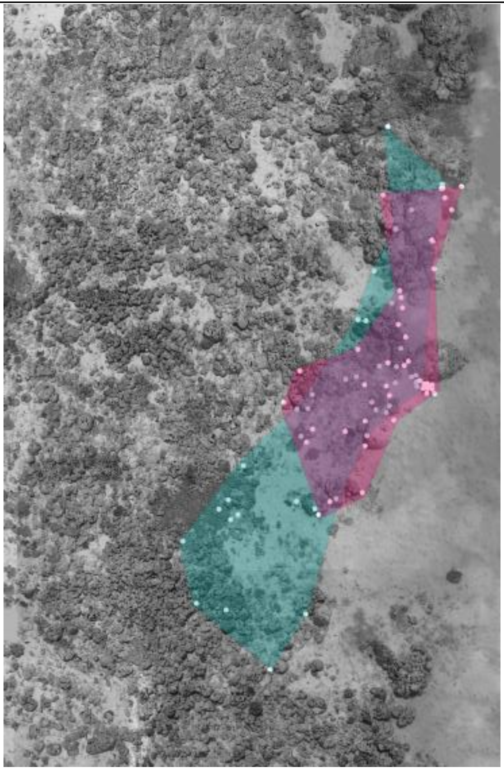 | 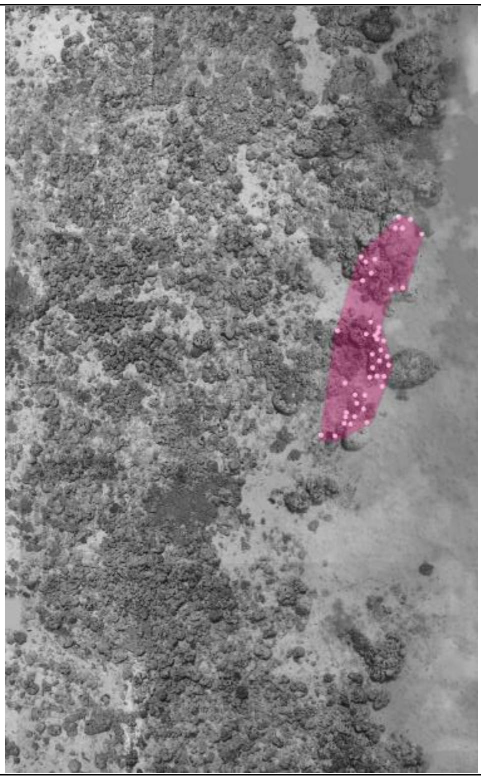 |

| **S14.2s : ORFPP** | |
|---|---|
| 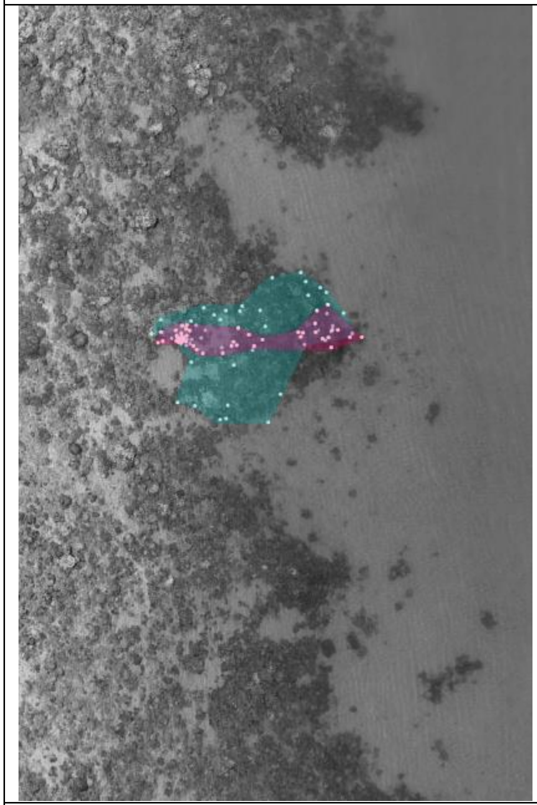 | 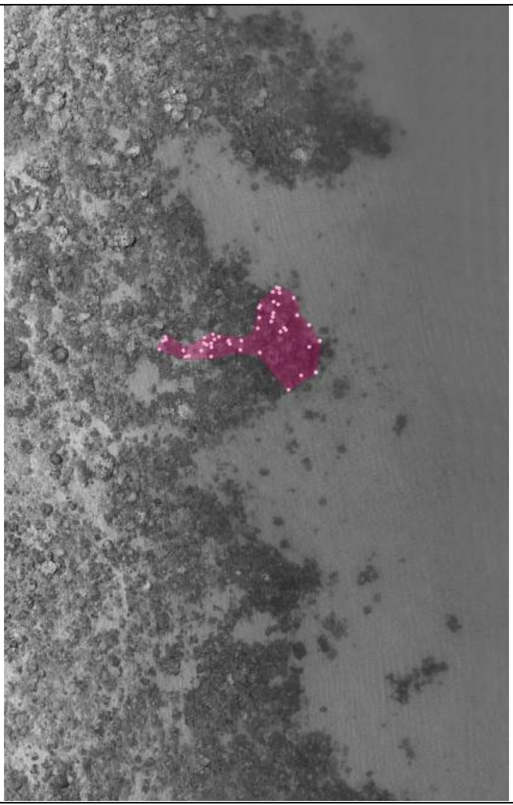 |
| **S14.2r : ORFGGMP** | |
| 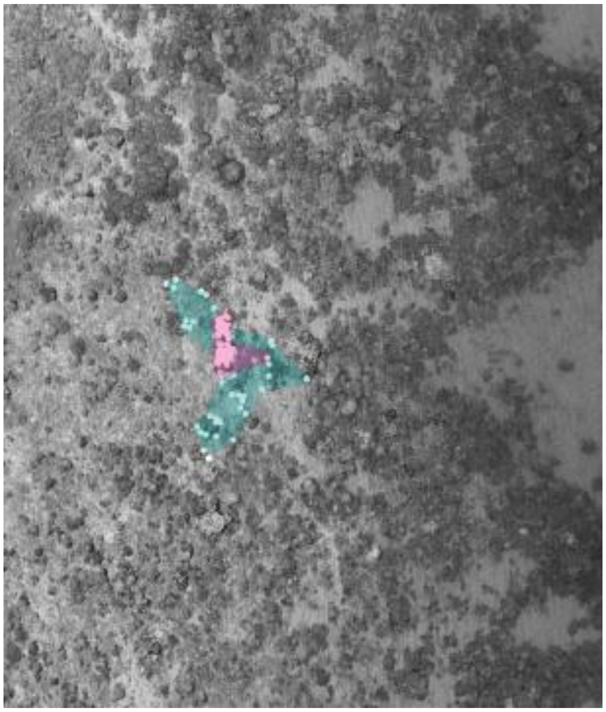 | 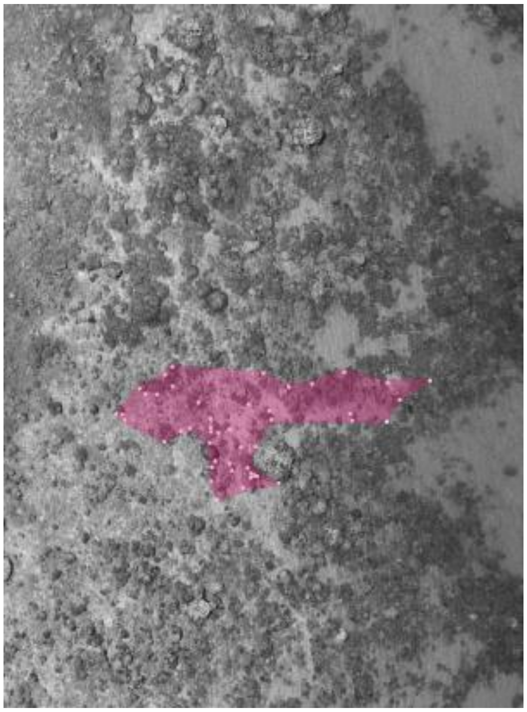 |

| S14.2q : OLMPY | |
| --- | --- |
| 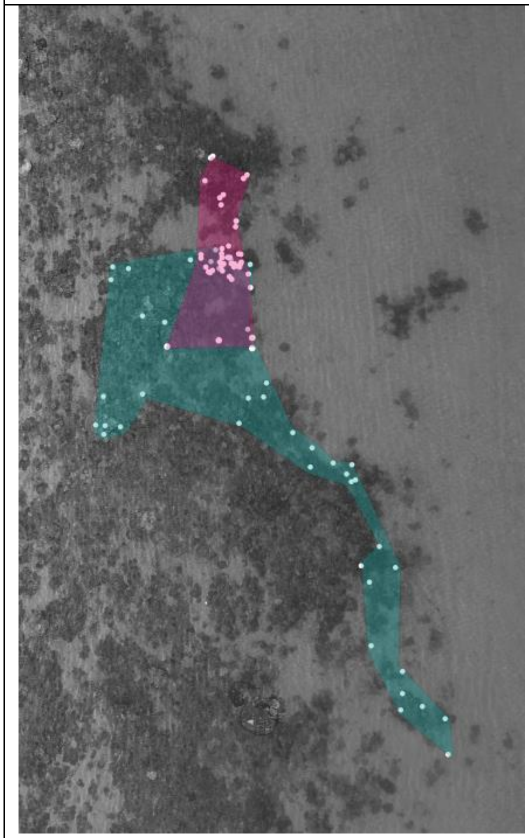 | 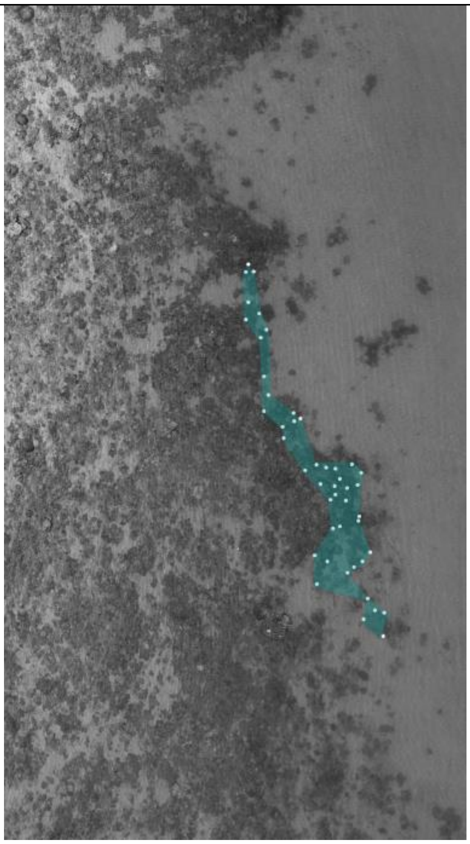 |
| **S14.2n : MLApprentist** | |
| 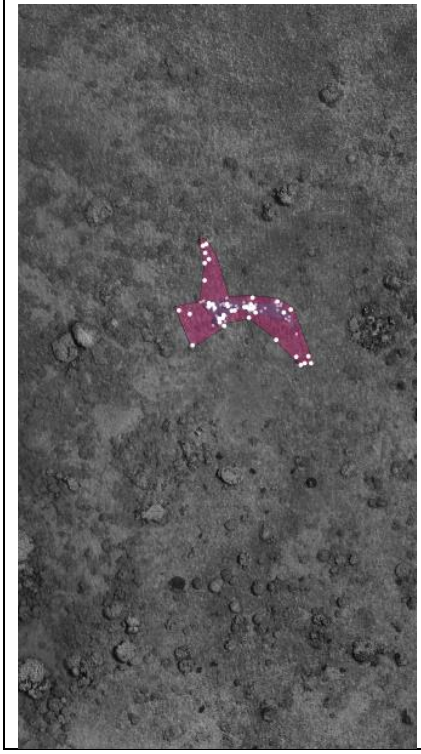 | 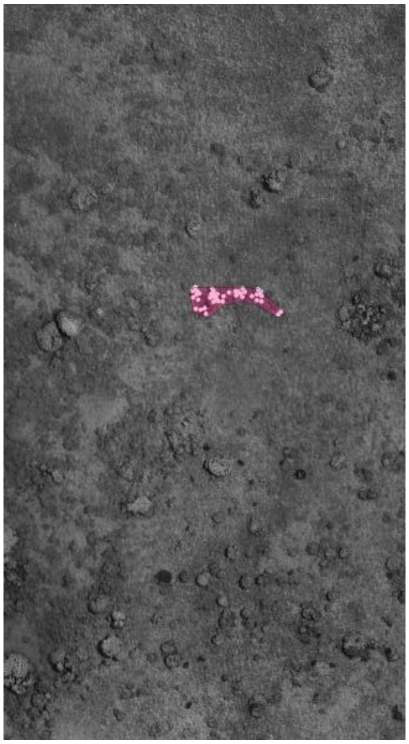 |

| **S14.2u : WLFPY** | |
|---|---|
| 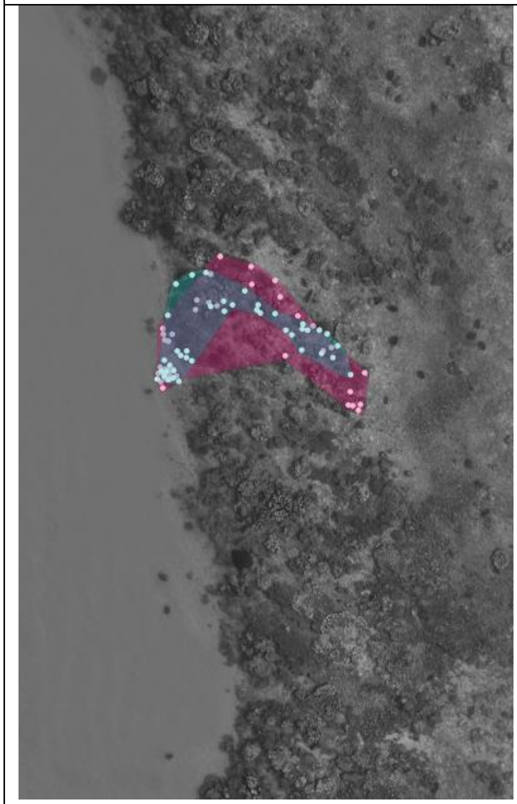 | 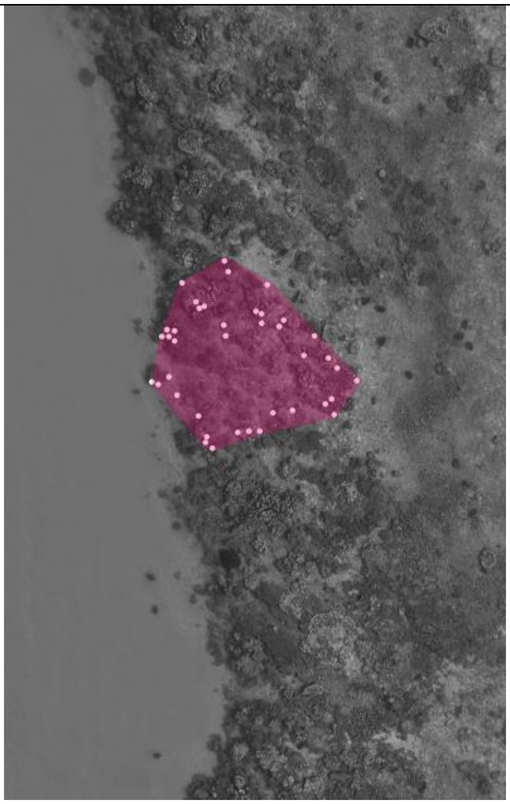 |
| **S14.2l : MBWierdo, MBIslGirl** | |
| 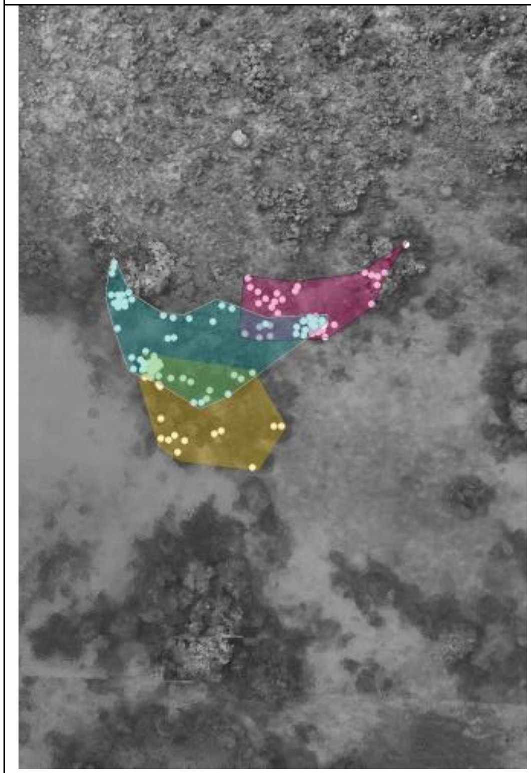 | 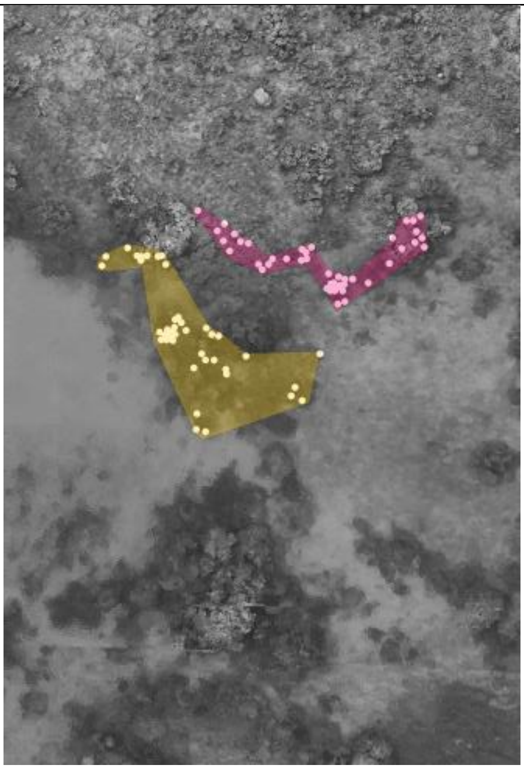 |

| S14.2a : CGLFGG | |
| --- | --- |
| 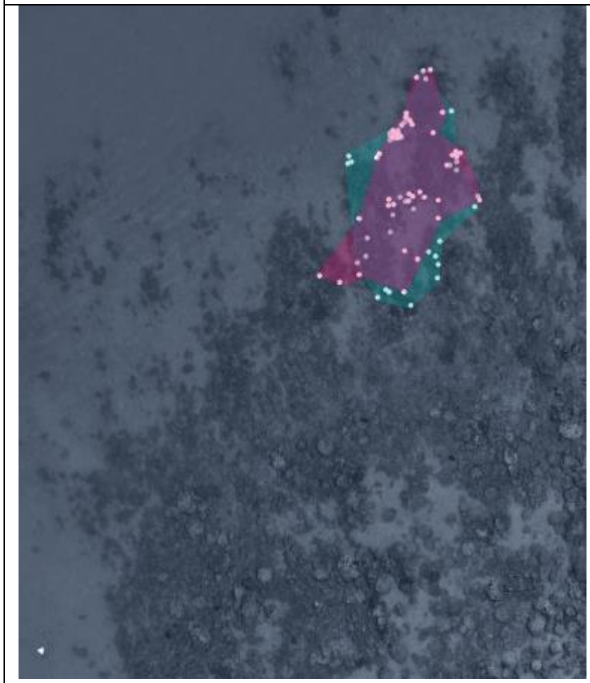 | 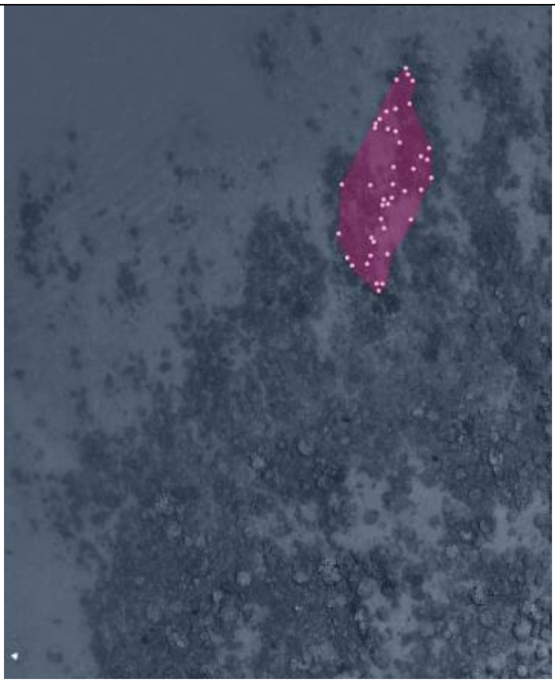 |

| S14.2i : LUBlackSpot2 | |
| --- | --- |
| 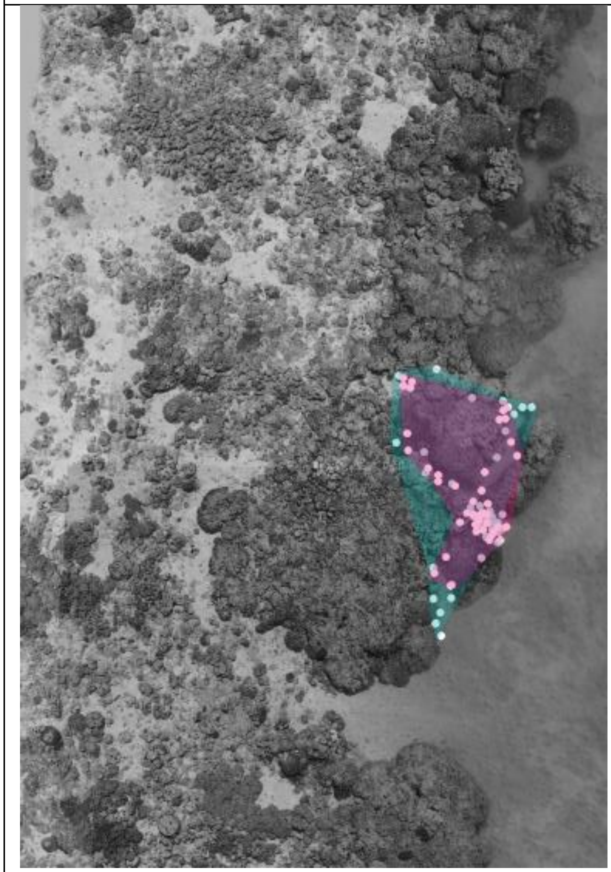 | 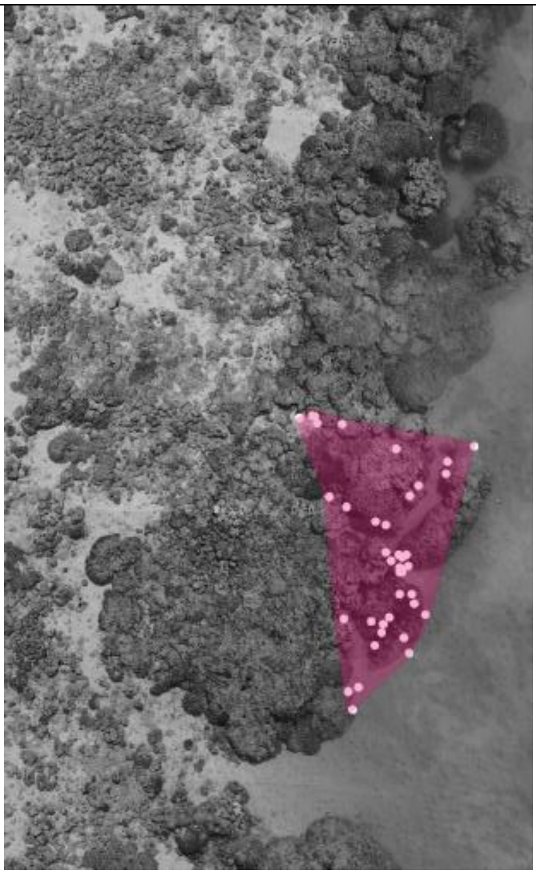 |

**S14.2k : MBB2Alpha**

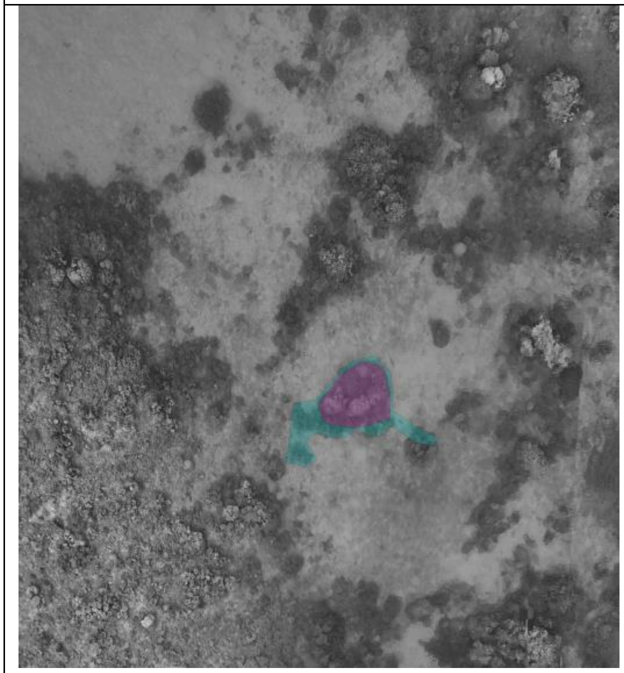

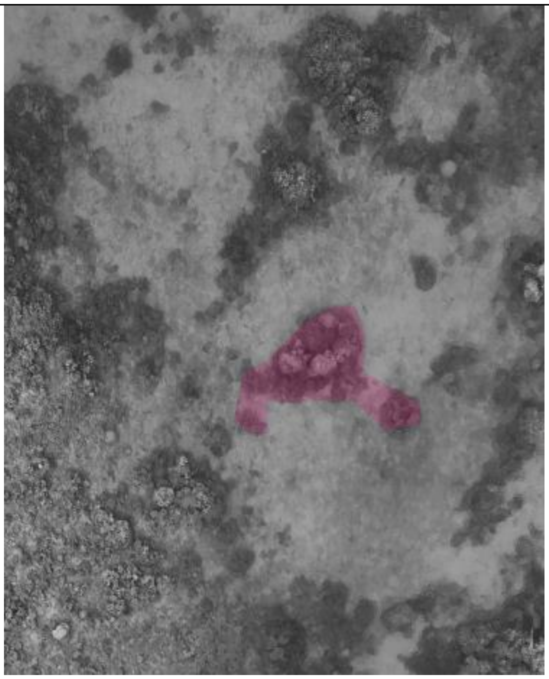

**S14.2o : MRAlphaRR, MRNewLFWW**

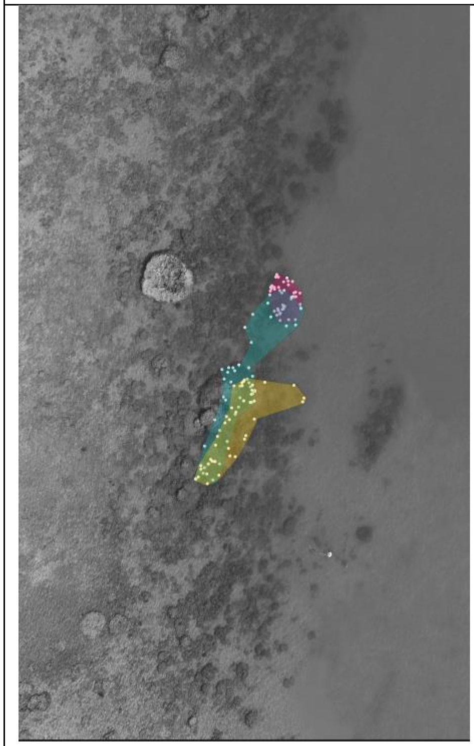

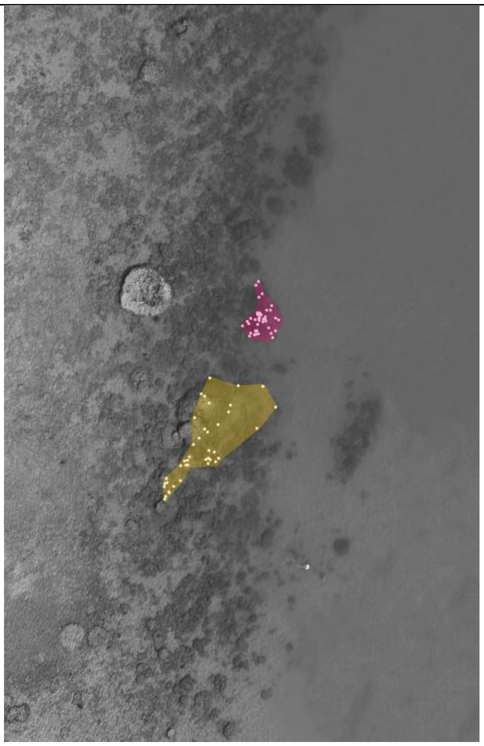

### Supplement 15: Migrant supplementary information

The following table details the number of juveniles and adult females (TL >= 50mm) gained by migrants after sex change.

#### Table S15: Migrants information

| Migrant | Sex Change Size | Nb Juveniles | Nb Adults | Social System |
|---|---|---|---|---|
| CGLFGG | 75.5 | 1 | 1 | Linear |
| CGNewLFPP | 77.9 | 1 | 4 | Linear |
| HLMRR | 59.7 | 1 | 0 | Linear |
| HRMYY | 71 | 0 | 2 | Linear |
| LUAnnFriend | 80.9 | 1 | 1 | Linear |
| LUBS2 | 89 | 0 | 5 | Linear |
| MLApprentist | 69.1 | 1 | 3 | Linear |
| MLApprentist2 | 75 | 0 | 1 | Linear |
| MLLFYR | 65.5 | 2 | 0 | Linear |
| MLNewRFWW | 77 | 4 | 5 | Branching |
| MRLMWW | 77.7 | 2 | 2 | Branching |
| MRNewLFWW | 77.6 | 2 | 3 | Branching |
| ONewBBFemale | 73.9 | 3 | 3 | Branching |
| ORFPY | 68.9 | 3 | 0 | Branching |
| WLFPY | 72 | 1 | 1 | Linear |
| LUAnninuccia | 93.7 | 1 | 1 | Linear |
